# Benchmarking computational methods for single-cell chromatin data analysis

**DOI:** 10.1101/2023.08.04.552046

**Authors:** Siyuan Luo, Pierre-Luc Germain, Mark D. Robinson, Ferdinand von Meyenn

**Author notes:** Correspondence to Ferdinand von Meyenn and Mark D. Robinson.

## Abstract

Single-cell chromatin accessibility assays, such as scATAC-seq, are increasingly employed in individual and joint multi-omic profiling of single cells. As the accumulation of scATAC-seq and multi-omics datasets continue, challenges in analyzing such sparse, noisy, and high-dimensional data become pressing. Specifically, one challenge relates to optimizing the processing of chromatin-level measurements and efficiently extracting information to discern cellular heterogeneity. This is of critical importance, since the identification of cell types is a fundamental step in current single-cell data analysis practices.

We benchmarked 8 feature engineering pipelines derived from 5 recent methods to assess their ability to discover and discriminate cell types. By using 10 metrics calculated at the cell embedding, shared nearest neighbor graph, or partition levels, we evaluated the performance of each method at different data processing stages. This comprehensive approach allowed us to thoroughly understand the strengths and weaknesses of each method and the influence of parameter selection.

Our analysis provides guidelines for choosing analysis methods for different datasets. Overall, feature aggregation, SnapATAC, and SnapATAC2 outperform latent semantic indexing-based methods. For datasets with complex cell-type structures, SnapATAC and SnapATAC2 are preferred. With large datasets, SnapATAC2 and ArchR are most scalable.

## Background

Recent advances in single-cell sequencing technologies have enabled the profiling of genome-wide chromatin accessibility and histone modifications, and allowed the exploration of epigenetic landscapes within complex tissues. However, the analysis of single-cell chromatin data is challenging due to two main reasons. Firstly, state-of-the-art technologies such as single-cell ATAC-seq (scATAC-seq) ^[1,2]^ and single-cell CUT&Tag (scCUT&Tag) ^[3]^ are based on DNA tagmentation, which produces sparse and noisy signals due to the low copy numbers and rare tagmentation events. It has been estimated that only 1–10% of accessible regions are detected per cell compared to corresponding bulk experiments ^[4]^. Secondly, unlike in single-cell RNA-seq data, there are no fixed feature sets for chromatin data. Usually, a set of genomic regions (e.g. bins or peaks) is first determined, and then the tagmentation events are counted within each region. For large genomes such as human and mouse, this leads to very high-dimensional data that not only raises challenges on the time and memory efficiency of the processing pipelines, but also hinders the statistical analysis. On the other hand, it is commonly assumed that single-cell data is sampled from a cellular state space that is of much lower intrinsic dimensionality than the observed data ^[5,6,7]^. Therefore, it is necessary and important to learn a low-dimensional representation of the data before further analysis.

In the past few years, there have been many efforts on improving feature engineering and dimensional reduction methods for scATAC-seq data. One idea is to use approaches that are originally designed for sparse and high-dimensional data (e.g. Latent Semantic Indexing and Latent Dirichlet Allocation from the natural language processing field), and directly apply them to the cell-by-region count matrix. Several popular methods fall into this broad category, although the underlying algorithms differ. For example, Signac ^[8]^ uses Latent Semantic Indexing (LSI), which is a linear dimensional reduction method consisting of a normalization step (e.g. Term Frequency - Inverse Document Frequency, TF-IDF) and Singular Value Decomposition (SVD); ArchR ^[9]^ employs an iterative procedure of LSI, in order to refine the feature selection during each iteration; cisTopic leverages Latent Dirichlet Allocation (LDA), a topic modeling method, to discern thematic structures; SnapATAC ^[10]^ uses diffusion maps, and SnapATAC2 uses Laplacian eigenmaps, both of which are non-linear dimensional reduction methods that work by constructing a graph representation of the data and then utilizing the eigendecomposition of some form of graph matrix. Another group of approaches first uses domain knowledge to aggregate the genomic region set into a much smaller set of meta-features such as motif hits, k-mers, genes, etc., and then applies dimensional reduction methods such as PCA on the cell-by-meta-feature matrix. For example, BROCKMAN ^[11]^ uses gapped k-mer frequency of the DNA sequence around insertion points, SCRAT ^[12]^ allows the usage of motifs, DNase I hypersensitive site clusters, genes or gene sets as features, and Cicero ^[13]^ calculates gene activity scores. A third type of method uses neural network models, such as PeakVI ^[14]^, which uses a variational autoencoder, and scBasset ^[15]^, which uses a convolutional neural network. Other ideas include integrating DNA sequence information, such as in scBasset and CellSpace ^[16]^.

Despite a large amount of available methods, there is currently no consensus on the best usage of these methods for scATAC-seq data. Chen *et al*. ^[4]^ did a benchmark on 10 methods and showed that SnapATAC, cisTopic, and Cusanovich2018 ^[17]^ outperform other aggregation-based methods. Since then, many new methods have been proposed ^[8,9,14,15,16]^, and an updated benchmark is desirable. Although a subset of methods has been frequently benchmarked in papers of new methods using a few popular datasets, it is hard to find an agreement between these benchmarking efforts. Therefore, a comprehensive and neutral benchmark effort is desired ^[18]^ to give an unbiased perspective on how these methods perform on a large variety of datasets.

One way to evaluate the feature engineering and dimensional reduction methods is to combine them with unsupervised clustering with the aim to identify cell types or cell states, which is a fundamental step for many downstream analysis ^[19]^. Previous benchmarking studies ^[4]^ for scATAC-seq data have focused on comparing the clustering outcomes at a single predefined resolution ^[20]^. However, determining the true number of clusters in advance is not always feasible, and as dataset complexity increases, the choice of clustering resolution becomes dependent on user-defined parameters and biological questions ^[21]^. Given that alterations in the number of clusters can have a substantial influence on many evaluation metrics ^[22,23]^, such evaluations may not fully capture the scenarios encountered by users during dataset processing and interpretation.

To provide a comprehensive assessment of the methods under investigation, we conducted our evaluation across three distinct levels: cell embeddings, graph structure, and final partitions. We employed a set of ten metrics to evaluate performance at each of these levels. By considering multiple aspects of clustering quality, our evaluation approach aims to provide a more thorough understanding of the strengths and limitations of each method. Based on our results, we provide guidelines for choosing analysis methods for different data types. Meanwhile, our data and analysis also provide a comprehensive framework for benchmarking common single-cell chromatin data analysis steps.

## Results

### Benchmark Design

To get a comprehensive understanding of the method performance, we used 6 published datasets of divergent sizes and sequencing protocols, and from different tissues and species (Table 1). In the absence of perfect ground truth, we included datasets with annotations from different information sources, including RNA modalities, genotypes, FACS-sorting labels, or tissue of origins. This ensures that our evaluation is not biased by specific assumptions of the ground truth. The coverage and signal-to-noise ratio (measured by transcription start site enrichment score, TSSE) also vary a lot across datasets, suggesting that our data collection represents a wide range of realistic test cases from different experimental protocols.

**Table 1:** Dataset overview.

| Dataset | #cells | Mdn TSSE | Mdn #frag | #classes | Annotation | Protocol | Species | Tissue |
| --- | --- | --- | --- | --- | --- | --- | --- | --- |
| Cell line <sup>[9]</sup> | 18538 | 1.449 | 4538 | 10 | genotype | 10X | human | cell line |
| Atlas1 <sup>[24]</sup> | 20204 | 14.049 | 3880 | 13 | tissue | sci-ATAC-seq3 | human | multiple |
| Atlas2 <sup>[24]</sup> | 15301 | 13.294 | 2939 | 10 | tissue | sci-ATAC-seq3 | human | multiple |
| 10XPBMC | 8560 | 13.595 | 13396 | 15 | RNA | 10X multiome | human | PBMC |
| Buenrostro2018 <sup>[25]</sup> | 1711 | 15.493 | 8556 | 9 | FACS labels |  | human | bone marrow |
| Chen2019 <sup>[26]</sup> | 5199 | 14.356 | 1355 | 13 | RNA | SNARE-seq | mouse | brain |

The benchmarking pipeline (Figure 1) starts from quality control (QC) and preprocessing to get the fragment files in BED format. These files serve as the input for each method. Then feature engineering and dimensional reduction are performed, and a cell embedding matrix is generated. This particular stage is where the various methods are applied. Subsequently, each embedding matrix is loaded into a common clustering and evaluation pipeline to get the clustering results.

**Figure 1:**
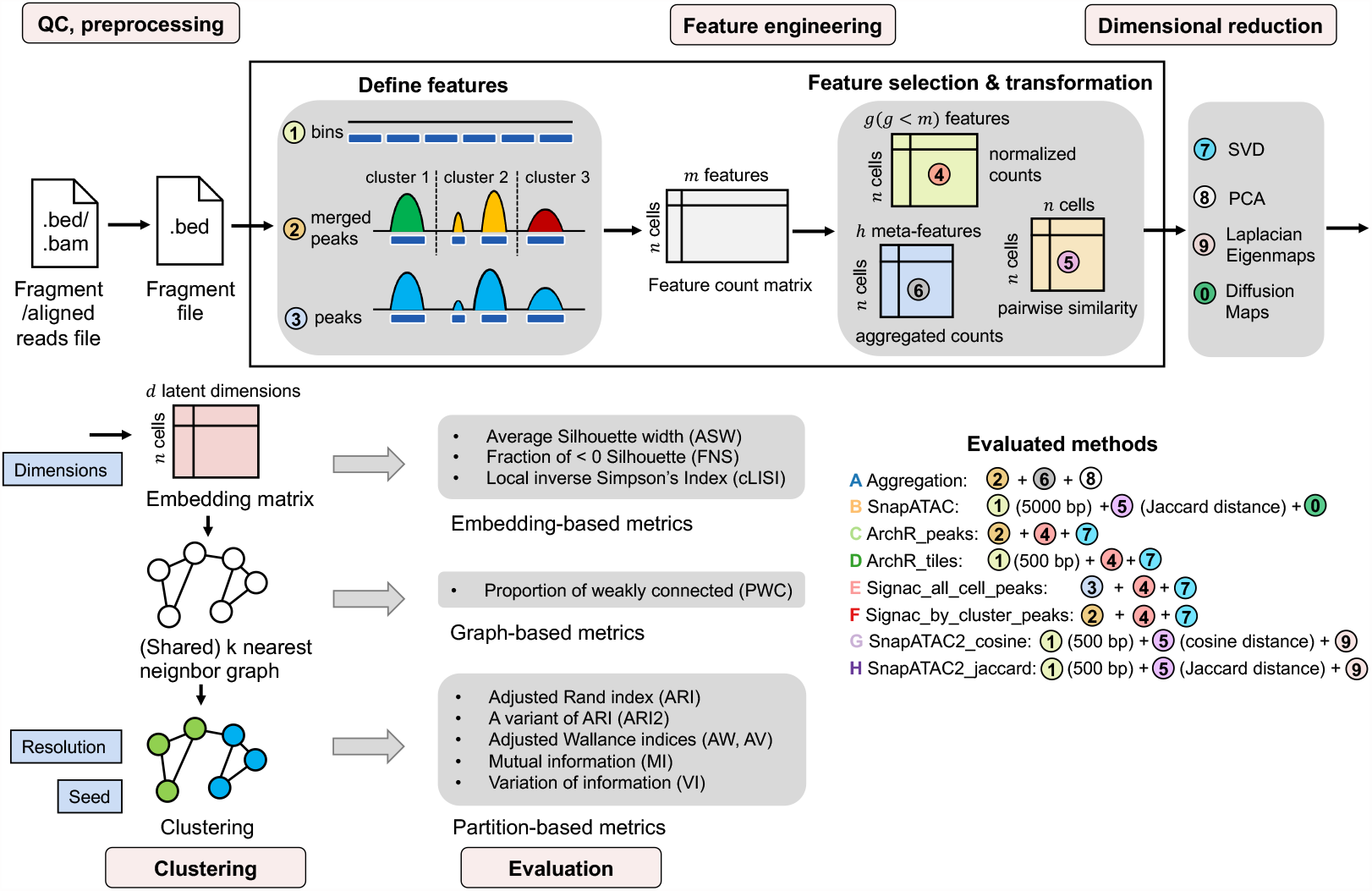
Benchmark framework. Starting from the .bed format fragment files or the .bam format aligned reads files, barcode-level QC is performed, and the filtered fragment files are input to the benchmark pipeline. The feature engineering step consists of two stages: (i) defining the genomic features; and, (ii) feature selection and/or transformation. Approaches for each stage are listed, and respective approaches of each method are indicated. Next, dimensional reduction is performed to generate the cell embedding matrix. A shared nearest neighbor (SNN) graph is constructed from the embedding matrix and then used for Leiden clustering. Evaluations are conducted on the embedding matrix, SNN graph, and final partitions, each using different metrics. During the evaluation, we explored multiple values for parameters such as the number of latent dimensions, resolution, and random seed - their positions within the workflow are denoted by blue boxes.

At the feature engineering and dimensional reduction stage, we benchmarked 5 methods in 8 configurations (Figure 1). Signac was included as a representative of LSI-based methods, and it uses a dataset-specific peak set as the genomic regions. Two different ways of defining this peak set were tested: (1) aggregate all cells for peak calling, or (2) first do coarse cell clustering, then do peak calling per cluster and use the merged peak sets from all clusters. The iterative LSI in ArchR was also included and tested on either genomic bins or merged peaks. In addition, we assessed SnapATAC and its recently updated version, SnapATAC2. For SnapATAC2, when calculating the pairwise cellular similarity matrix, it allows the usage of either Jaccard or Cosine distance. So we tested both metrics to see how appropriate they are for the sparse and near-binary data. Besides, while aggregating regions based on biologically meaningful features such as motifs has tended to have poor performance ^[4]^, *ad hoc* feature clustering and summing into meta-features was shown to be a viable strategy for doublet detection ^[27]^. We therefore chose to include this strategy as well in our evaluation.

In the clustering process, several parameters could affect the clustering performance. We explored various values of these parameters to study their effects (Figure 1). These parameters include the number of latent dimensions, resolution, and random seed used in Leiden clustering. Meanwhile, during the evaluation process, we assessed each method at different clustering steps including the cell embedding, the shared nearest neighbor (SNN) graph, and the partition level, to eliminate the effect of potentially suboptimal parameter choice.

### Method performance is dependent on the intrinsic structure of datasets

Among our six datasets, Cell line, Atlas1, and Atlas2 consist of mixed cell lines or cell types from various tissues. These three datasets show a relatively simple structure, with distinct cell clusters and little hierarchy. Conversely, the remaining three datasets, derived from specific tissues, carry inherent complexity, including closely related subtypes and/or hierarchical structures. This division is reflected by the average levels of many evaluation metrics: the simpler datasets show a higher average ARI and lower cLISI and PWC score, and conversely (Figure 2, Figure 3a). Through our analysis, we noticed that some methods performed relatively better on the simpler or more complex tasks.

**Figure 2:**
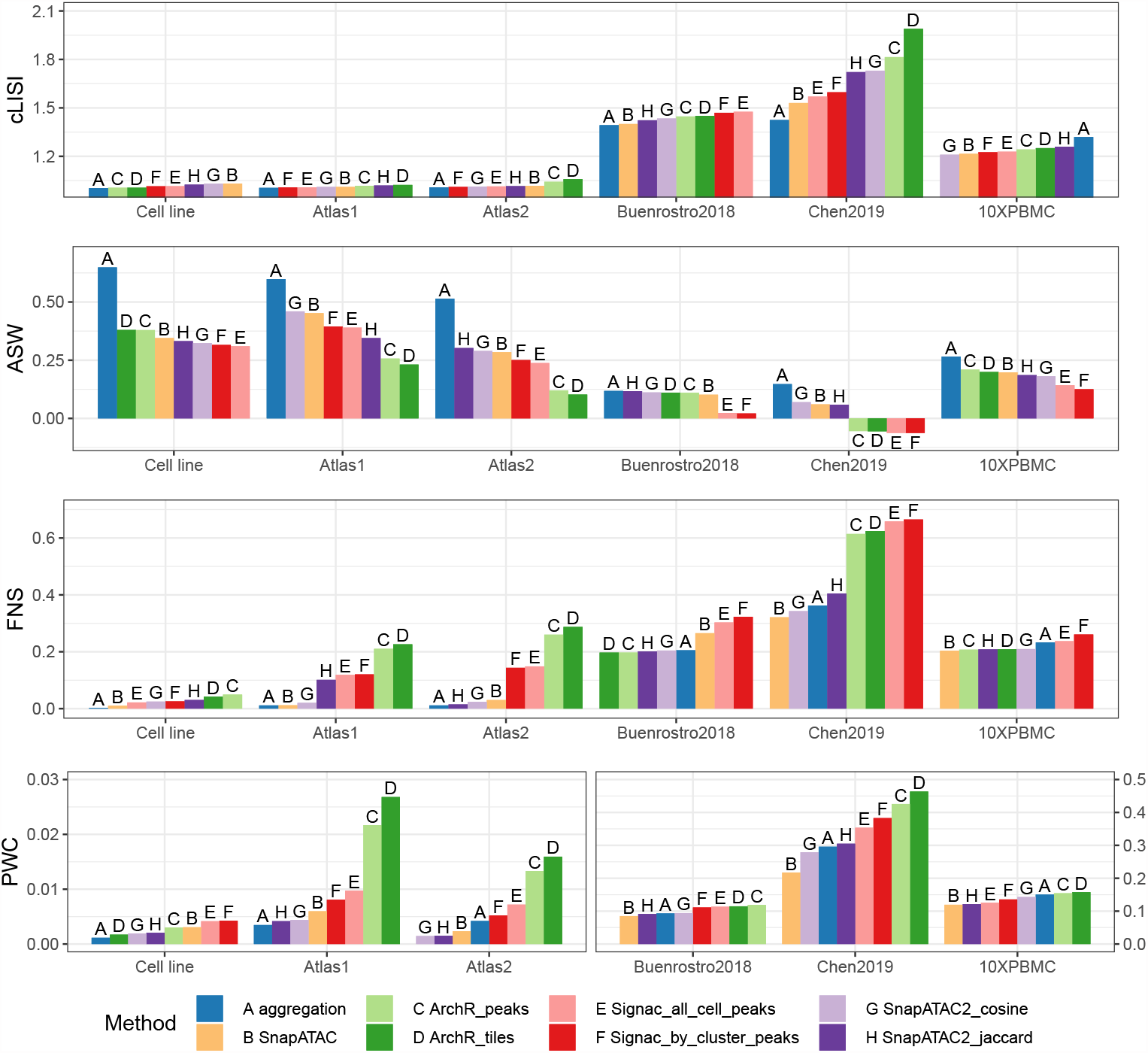
Embedding- and graph-level metrics averaged across all cells or all cell classes for each dataset; each subpanel represents an evaluation metric. Bars are sorted from the best to the worst performance.

**Figure 3:**
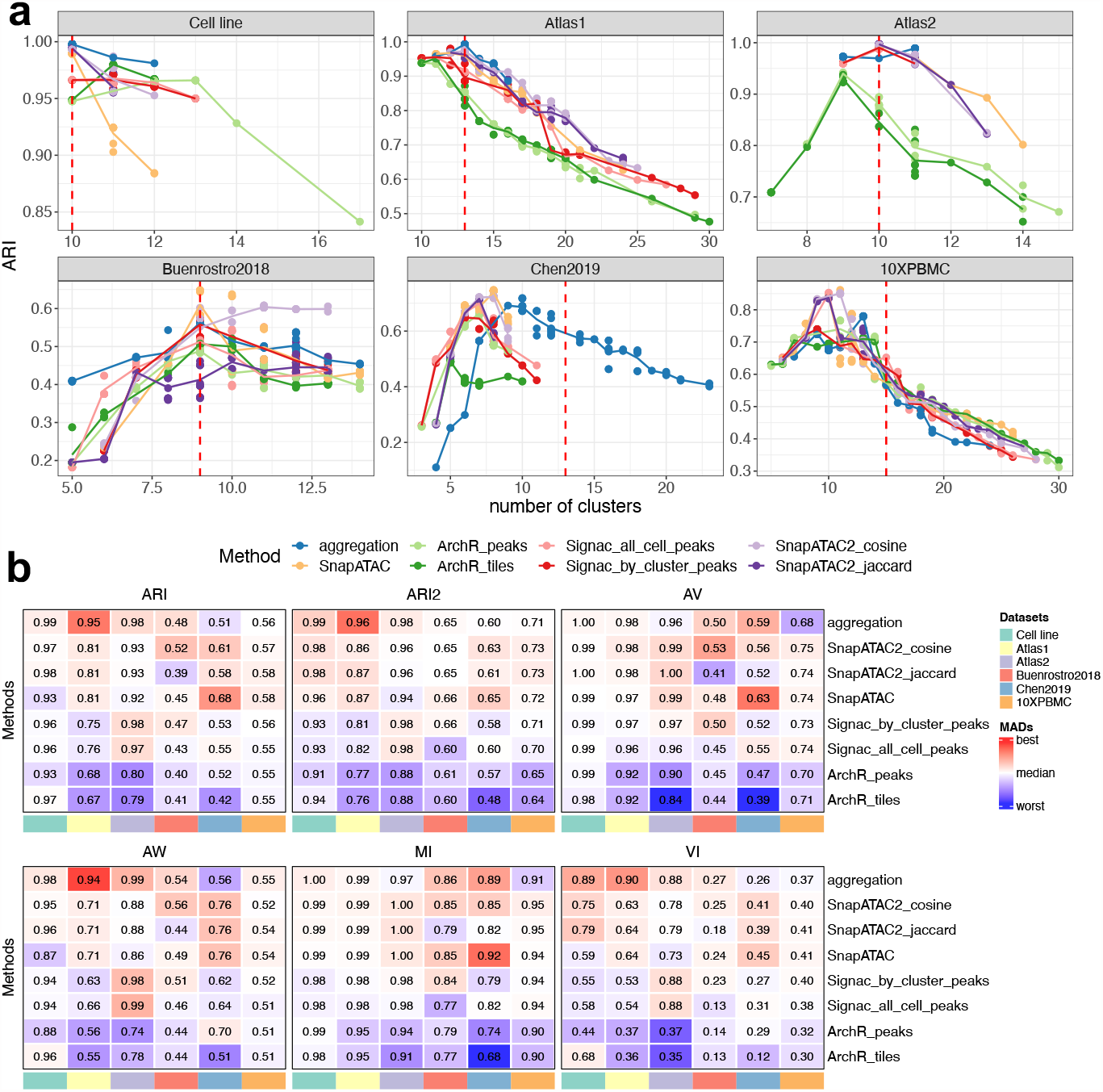
Clustering results. **a** The adjusted Rand Index (ARI) plotted against various number of clusters; each subpanel represents a dataset. Each point represents a clustering solution obtained by varying the resolution parameter and the random seed in Leiden algorithm. The line plot is the average ARI at a given number of clusters. **c** Heatmaps displaying the normalized areas under the curve (AUC) from plots similar to **a**, but for various partition-level metrics. The color scale indicates deviations from the column-wise median scaled by the matrix-wise median absolute deviation ^[23]^. It provides a comparison of relative performance between methods, and is unified across datasets and robust to outliers.

As mentioned above, we first evaluated the cell embedding and SNN graph-level outputs (Figure 2). The cluster LISI score (cLISI), which measures the purity of neighborhood composition in the embedding space, was always close to 1 in easy tasks and showed little discrimination between methods. This indicates that most local neighborhoods (*k* = 90) contain a single cell type in the embedding space. On the contrary, the Silhouette width is calculated between a cell and a whole cluster (*k >* 300). We observed that the average Silhouette width (ASW) sometimes showed inconsistent rankings compared to other metrics. While the Silhouette score has commonly been utilized for benchmarking clusterings in single-cell datasets, a potential issue is that Euclidean distance may not be suitable for accurate assessments in high-dimensional spaces over long-range distances ^[5]^. Therefore, we consider the Silhouette score calculated using Euclidean distance at the cluster level to be less appropriate for our analysis. Nevertheless, we considered the fraction of negative Silhouette (FNS) to be relatively more robust to certain space transformations and thus more appropriate here. In two of the three relatively easy tasks, ArchR peaks and ArchR tiles exhibited the highest FNS, followed by Signac all cell peaks and Signac by cluster peaks, while SnapATAC, SnapATAC2 cosine, and aggregation always displayed close to 0 FNS. Consistently, ArchR showed the worst PWC score in two of the three easy tasks, followed by Signac.

At the clustering level, the 6 evaluation metrics measure multiple aspects of the clustering performance (Figure 1). ARI, VI measure the overall agreement between the clustering results and the ground-truth annotation. ARI2 adjusts for the class size bias and is more sensitive to errors in small classes. AV and MI reflect mostly the homogeneity of clusters (i.e. the degree to which it includes only cells of one class), while AW represents the completeness of true classes in the clustering. Since the clustering solution varies by using different resolutions and random seeds, these metrics would also vary across these parameters. As shown in Figure 3a, ARI is strongly affected by the number of clusters. Usually the best ARI is achieved at the cluster number that is equal or close to the ground-truth number of classes, and then as the cluster number increase or decrease, ARI can deteriorate dramatically. We therefore inspected multiple combinations of resolutions and random seeds that give different numbers of clusters, and summarized the results using normalized areas under the curve (AUC, see Methods), as shown in Figure 3b. The AUC of ARI showed that ArchR related methods tend to have a lower rank than SnapATAC2 based methods, and that datasets 10XPBMC showed less discrimination between methods than other datasets, which is consistent with Figure 3a. These observations confirm the use of AUC as a good summary of results across parameters.

For clustering tasks that are relatively easy, the number of clusters that provides the highest ARI value is usually equal to the number of classes of the ground truth (Figure 3a), with a few exceptions. ArchR tiles and ArchR peaks achieved the best ARI using fewer clusters than the true classes in datasets Atlas1 and Atlas2, and then their performance starts to deteriorate as the number of clusters increases. This is because ArchR failed in separating small similar classes from one another, and instead segregated other classes (Figure 4a,b, Figure S2a,b,c,e). Signac all cell peaks and Signac by cluster peaks are among the best performing methods in the Atlas2 dataset (Figure 3a,b), but also segregated classes in Atlas1 (Figure S2f,g). In contrast, SnapATAC, SnapATAC2 cosine, SnapATAC2 jaccard, and the aggregation method performed consistently well for the easy tasks in the sense that the optimal ARI is always achieved at the correct number of clusters (Figure 3a); in addition, the clustering is nearly perfect (Figure 4c,d). As noted in the cell line task, SnapATAC seems to have the worst performance according to ARI when over-clustering (Figure 3a), but this is because it segregated a large class (293T) compared to what SnapATAC2 segregated (GM12878) (Figure S1b,c); this is an example of the cluster size bias of ARI. The adjusted version, ARI2, does not show a preference for SnapATAC2 over SnapATAC (Figure S1a), further underscoring the importance of considering multiple evaluation metrics. Other partition-based metrics indicate that aggregation, SnapATAC, and SnapATAC2 are the best methods, followed by Signac; ArchR performed worst (Figure 3b). Overall, for easy tasks, SnapATAC, SnapATAC2, and the aggregation method performed the best, followed by Signac, while ArchR had a difficult time correctly clustering rare cell types.

**Figure 4:**
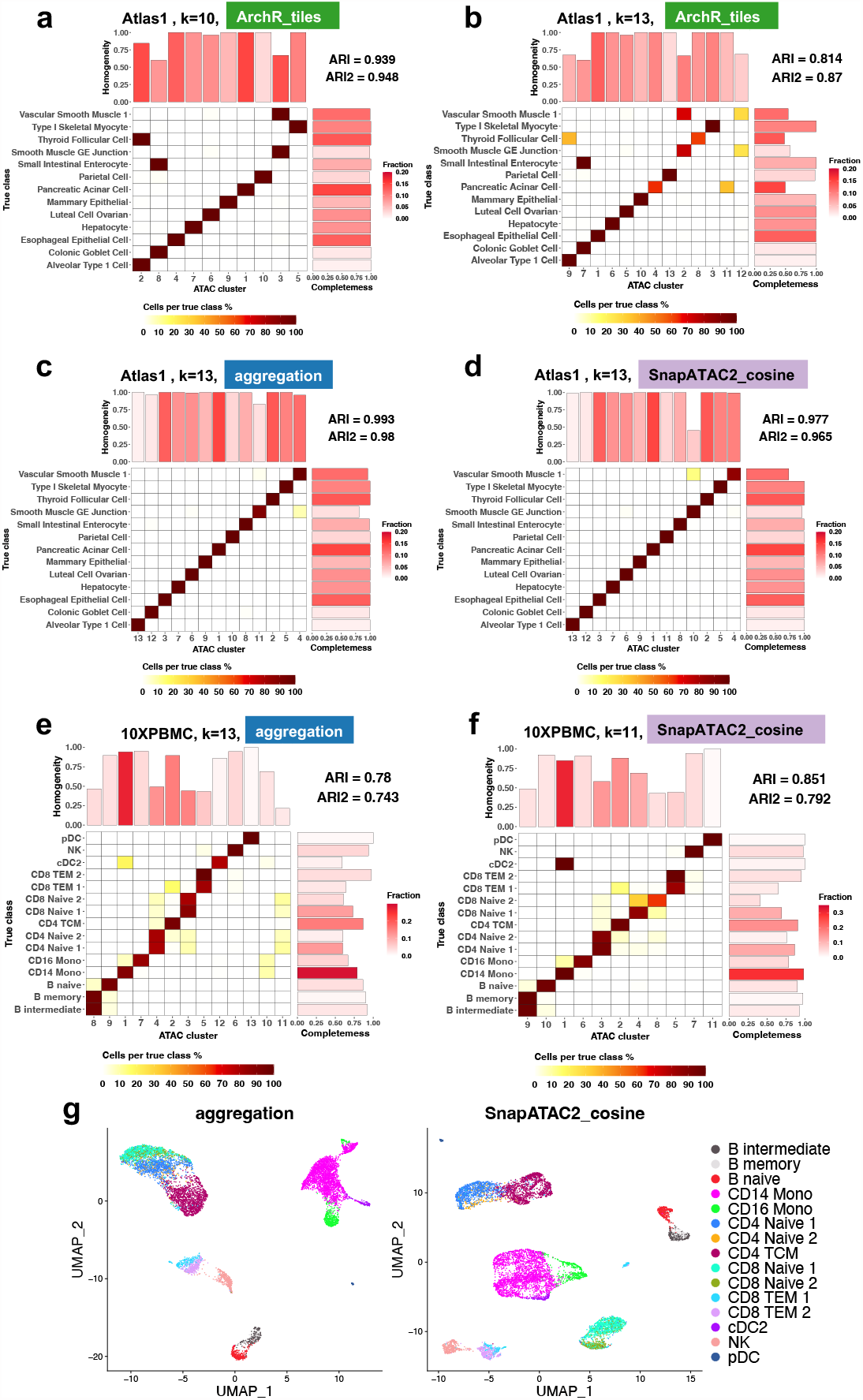
**a-f** True classes and their fractions of agreement with the predicted clusters. **a-b** are ArchR tiles on Atlas1, **c-d** are aggregation and SnapATAC2 cosine on Atlas1, and **e-f** are aggregation and SnapATAC2 cosine on 10XPBMC. The x-axis is the predicted clusters, and the y-axis is the ground truth classes. The colors of tiles indicate the proportion of cells from the corresponding true class (each row sums up to one). A clearer diagonal structure indicates better agreement. ARI and ARI2 are calculated and shown on the top right. The barplot on top shows the value of AV (Methods) and can be interpreted as the homogeneity of the corresponding clusters. The barplot on the right shows the value of AW and represents the completeness of each true class in the prediction. The color of the bars shows the proportion of cells in each cluster/ground truth class. In title, the corresponding datasets, methods, and number of clusters are indicated. **g** Corresponding UMAP given by aggregation and SnapATAC2 cosine on dataset 10XPBMC. The aggregation method did not resolve correctly CD14 vs CD16 monocytes, as well as CD4 vs CD8 naive T cells.

For complex datasets, the overall cLISI, FNS, and PWC increased compared to simple datasets (Figure 2). Specifically, on average more than 20% cells of each true class have a negative Silhouette score, and more than 8% cells of each true class are weakly connected to the belonging communities in the SNN graph. In datasets Buenrostro2018 and 10XPBMC, cLISI and PWC did not show much discrimination between methods. FNS is also similar across methods in 10XPBMC, although slightly worse for Signac all cell peaks, Signac by cluster peaks, and SnapATAC in Buenrostro2018. In Chen2019, the aggregation method exhibited the best local neighborhood purity reflected in the cLISI score, followed by SnapATAC, Signac all cell peaks, and Signac by cluster peaks, while ArchR tiles and ArchR peaks were the worst. FNS indicated that for Signac and ArchR, more than 60% cells of each true class have a negative Silhouette, while for other methods, this value is less than 40%. In alignment with this, ArchR and Signac also exhibited the worst PWC values in this dataset.

When comparing the clustering results, we noted that the number of clusters of the highest ARI does not always equal the number of classes in the annotation for difficult clustering tasks (Figure 3a). This appears to be because of populations that are hard to separate, either because they are rare, as in the Chen2019 dataset (Figure S3), or because they are very similar to each other, as intermediate and memory B cells in the 10XPBMC dataset (Figure 4e,f,g).

For these difficult clustering tasks, SnapATAC and SnapATAC2 cosine consistently performed the best, while ArchR tiles and ArchR peaks were the worst (Figure 3a,b). SnapATAC2 jaccard seems to have a worse performance than SnapATAC and SnapATAC2 cosine in Buenrostro2018 according to ARI and AV, but it showed a comparative performance of ARI2. The aggregation method also showed good performance in Buenrostro2018 and Chen2019, but slightly underper-formed in 10XPBMC. The decrease in performance of aggregation in 10XPBMC is mostly because of the mixing of subtypes, as AV and MI indicate (Figure 3b), and is clear in Figure 4e,g. Signac tended to perform better than ArchR no matter which configuration was used, and in dataset Buenrostro2018 it was comparable to SnapATAC and SnapATAC2. Overall, for difficult tasks, we found that SnapATAC, SnapATAC2 cosine, and SnapATAC2 jaccard are the top methods, followed by the aggregation approach, then Signac, while ArchR is the worst.

### Method choices at different feature engineering steps

This section focuses on the analysis of various choices made during the process of feature engineering, specifically regarding genomic features, peak calling methods, and distance metrics, and their impact on the overall clustering performance.

### Peaks versus bins

Due to the absence of a standard feature set for chromatin data, researchers often resort to using either genomic bins or peaks, each with its own limitations ^[28]^. Genomic bins suffer from the arbitrary selection of bin length and break-up positions. On the other hand, peaks align more closely with functional intervals, but present challenges in their identification in rare cells, and require additional processing when integrating different datasets.

To assess the clustering performance with these two types of genomic features, we compared the results obtained using ArchR tiles (non-overlapping genomic bins of 500bp) and ArchR peaks (MACS-2 consensus peaks across clusters) (Figure S5a, Figure S6a). Among most datasets we inspected, the performance of these two approaches is very similar across our metrics. This is consistent with claims in the literature ^[29]^. Only in the Chen2019 dataset, ArchR peaks exhibited a slightly better cLISI score (Figure S5a) and higher clustering-level performance than ArchR tiles (Figure S6a), mostly because it separated classes “L4 1” and “L4 2” better (Figure S3). In the Atlas2 dataset, the utilization of peaks demonstrated improved cluster homogeneity, as evidenced by metrics such as AV, MI, and FNS, but not necessarily enhanced class completeness in AW (Figure S5a, Figure S6a). In summary, we found that the use of peaks exhibited marginal or no significant improvement over the use of bins.

### Peak calling methods

If peaks are used as the genomic features, to facilitate the identification of population-specific peak sets, a common approach is to employ a two-step peak calling procedure, in which cells are first clustered using global peaks, before a second round of per-cluster peak calls. In our study, we tested two Signac pipelines, namely Signac all cell peaks and Signac by cluster peaks, to compare the effectiveness of one-step versus two-step peak calling. We observed that at the embedding and graph level (Figure S5b), these two approaches showed nearly identical performance. At the final partition level, the performance was still very similar in easy tasks, and only on specific difficult datasets did one method perform slightly better than the other. Specifically, in the Buenrostro2018 dataset, Signac by cluster peaks outperformed Signac all cell peaks (Figure S6b). Whereas in the Chen2019 dataset, Signac all cell peaks yielded slightly better results (Figure S6b, Figure S4a,b), mostly because L6 IT and L5/6 IT cells are not properly grouped by Signac by cluster peaks. Overall, our findings revealed that the two-step approach does not always outperform the one-step approach.

The two-step peak calling strategy somewhat mirrors the underlying concept of iterative LSI, where features of a subsequent round are dependent on previous clustering results. Considering that ArchR consistently underperformed in our benchmarks, we reason that this iterative approach might contribute more to stabilizing feature selection rather than significantly improving the discernment of difficult-to-distinguish cell types.

### Distance metrics

Despite the debate between using peaks or bins, scATAC-seq data is usually regarded as binary, and therefore in SnapATAC2, either Jaccard or cosine similarity was used to construct the affinity matrix. We observed in our results that both similarity metrics showed very similar cLISI and PWC scores across datasets; using cosine similarity gave better FNS scores in two of the six datasets (Figure S5c). Metrics of clustering results indicated that the performance of cosine similarity was very similar to Jaccard similarity (Figure S6c), especially after being adjusted for class size effects using ARI2. Overall, our results proved that both similarity metrics work comparably well in these clustering tasks.

### Dimensions of the latent space

The five methods we evaluated use different underlying algorithms for dimensional reduction. ArchR and Signac use truncated SVD, which identifies and preserves directions of maximum variance in the data. SnapATAC and SnapATAC2, on the other hand, apply graph-based spectral embedding. Specifically, SnapATAC2 uses Laplacian Eigenmaps, which removes higher-frequency variations from one node to its neighbors, and preserves low-frequency structures of the graph ^[30]^. In the aggregation method, feature-level aggregation is performed, followed by principal component analysis (PCA). The goal is to exploit the redundancy of the high-dimensional genomic feature space to average out potential noise.

Considering the varying assumptions and goals of each dimension reduction method, we investigated if different numbers of dimensions across methods could contain distinct information, and how the choice of the number of dimensions for the embedding space affects performance. We examined a series of *d* values, namely 15, 30, 50, 100, and calculated the embedding-level and graph-level evaluation metrics (see Figure S7 a). We observed that SnapATAC and SnapATAC2 were particularly sensitive to this parameter, with performance rapidly deteriorating as *d* increased. This trend is observable in the increasingly blurred structures in UMAP in Figure S7 b, and suggests that the later dimensions may contain less cell-type-relevant information. In contrast, the aggregation method demonstrated robustness to this parameter across most datasets. This aligns with the assumption that the aggregation method removed the noise by averaging it out, so that later dimensions are also smoothed signals. Signac and ArchR displayed an intermediate trend, and later eigenvectors may also have a smaller signal-to-noise ratio, especially in more complex datasets.

### Stability of clusterings and robustness of clustering performance

We observed that in certain cases, the inherent randomness of Leiden algorithm (Methods) can lead to instability in the clustering results (e.g. Figure S8). To account for this, we performed the clustering steps using 5 different random seeds, and compared the clustering results by calculating pairwise ARI (Figure S9 and Figure S10). The variability in the clustering outcomes varies depending on the datasets, methods, and resolution parameters employed. Generally, using the same resolution value yielded partitions with the same number of clusters. However, in some cases, changing the random seed resulted in partitions with different cluster numbers, leading to increased variability. Notably, in simple datasets, forcing over-clustering of cells by increasing resolution tends to amplify the variability (Figure S10). This can be attributed to the fact that the simpler datasets only have flat clustering structures, and increasing the resolution merely introduces random splits within the true communities.

We then focused on the resolution value that yielded the highest clustering performance, as measured by ARI against the ground truth. Specifically, we looked at how much the pairwise ARIs between seeds deviate from 1, which reflects the level of instability in the clustering outcomes. Interestingly, we found a positive correlation between the deviation and PWC value of the SNN graph (Figure S11 a). Furthermore, we observed that the variation in clustering outcomes was more prominent across datasets rather than between methods.

In order to assess the impact of clustering result variability on the evaluation of clustering performance, we examined the variability of performance measurements using different random seeds. Notably, despite the instability observed in the clustering results, we found that the similarity to the ground truth remained relatively consistent (Figure 5 a, coefficient of variation (CV) of ARI; Figure S12). This consistency provides us with a solid foundation for confidently interpreting the evaluation results.

**Figure 5:**
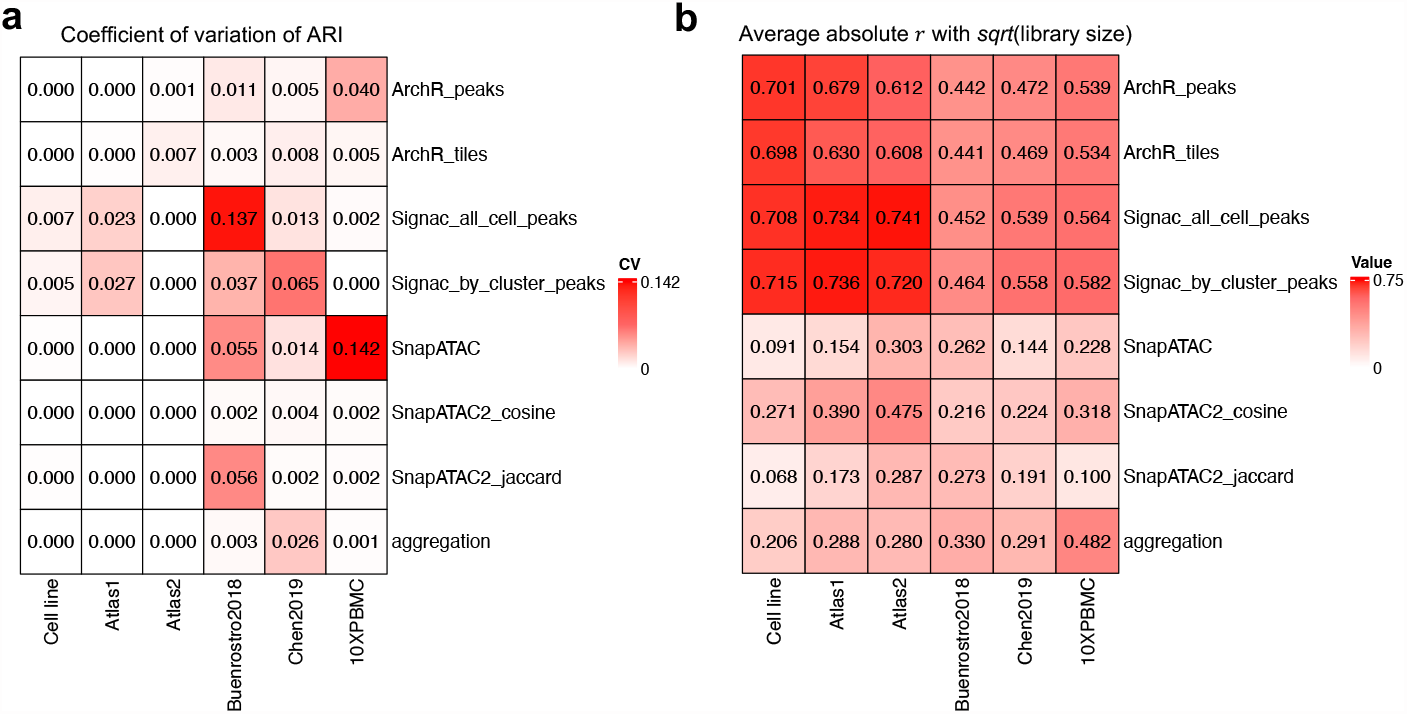
**a** Coefficient of variation of ARI between predicted clusterings and the true cell types. **b** Average absolute Pearson’s correlation with fragment counts; For a given latent dimension and a given cell type, the Pearson’s correlation coefficient is calculated between the latent axis and the square root of fragment counts. This correlation value was then averaged across cell types and across the first 10 latent components.

### LSI-based methods show strong library size biases

Large library size variation arising from technical biases are often observed in single-cell data, and can potentially confound the downstream analysis ^[19]^. Therefore, we examined to what extent the cell embedding of each method was driven by library size. By looking at the scatter plot of each latent dimension against the empirical library size, one can see that LSI-based methods (Signac, ArchR) showed a strong library size bias across all datasets (Figure S13). The difference between cell types may reflect biological variation, since global chromatin accessibility can differ during cell differentiation. However, the difference within each cell type is more likely due to technical aspects such as sampling effects. Therefore, we quantified this bias by calculating, for each latent dimension, the Pearson’s correlation coefficient *r* with the square root of library size per cell type, and averaged the absolute value across all cell types. This value is further averaged across the first 5 components (Figure 5 b, average absolute correlation). Note that we followed the suggestion in Signac’s tutorial and always removed latent components *r >* 0.75 with the library counts. In all our datasets, this criterion always removed the first component of Signac and ArchR embeddings, while no component was removed for other methods. After this filtering, the average absolute correlation is still between 0.5 to 0.75 in Signac and ArchR, followed by aggregation-based method and SnapATAC2 using cosine distance, where the correlation is around 0.2 - 0.5, then SnapATAC and SnapATAC2 using Jaccard distance, with correlation smaller than 0.3. We further evaluated how the library size bias affects the whole cell embedding by calculating the spatial autocorrelation of library size on the k nearest neighbor graph using Geary’s C index (Figure S11 b). Signac and ArchR tend to have a positive spatial autocorrelation, while aggregation, SnapATAC, and SnapATAC2 tend to have a smaller autocorrelation. In conclusion, LSI-based methods such as Signac and ArchR generate latent representations that are strongly associated with library size, while SnapATAC and SnapATAC2 using Jaccard distance are less affected.

### Time and memory complexity

Due to the large feature space in scATAC-seq data, it is crucial to use methods that scale efficiently in terms of time and memory usage. We monitored the CPU time and peak memory usage in our Snakemake pipeline (Figure 6 a,b). For the aggregation method, we tracked the program either from the start of peak count matrix generation (aggregation + Signac) or subsequent to it. We found that SnapATAC2 performed the best in terms of runtime, while ArchR is the most memory efficient. SnapATAC had low memory consumption with small datasets; however, its memory usage increased rapidly as the dataset size increase, making it the least scalable option.

**Figure 6:**
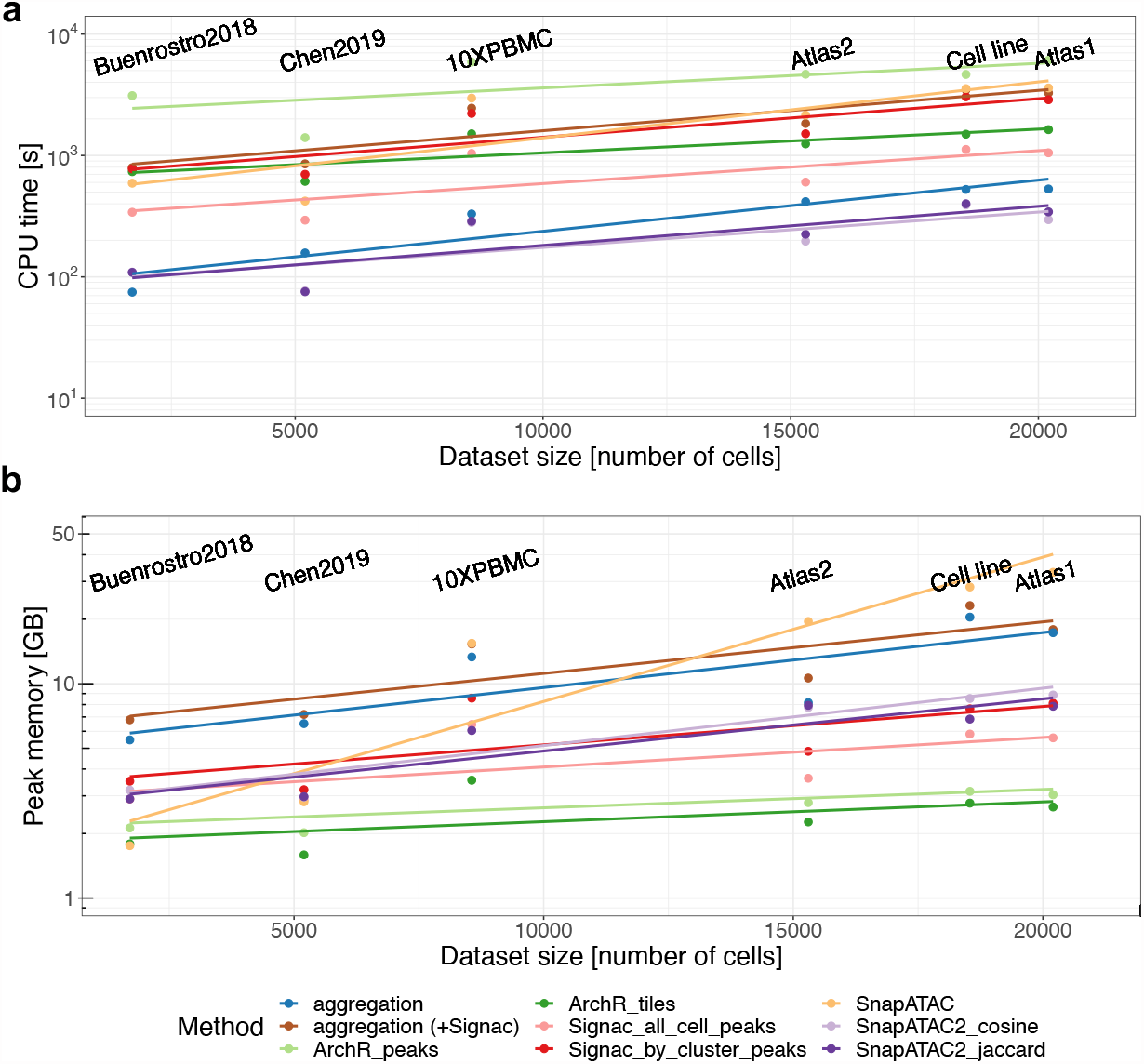
The CPU time and peak memory usage of each method across datasets of different sizes.

ArchR and SnapATAC2 both use on-disk storage instead of loading the entire dataset into memory. This is achieved by storing large-sized data in an HDF5-format file on disk and using an object to store small-sized metadata, which contains references to the corresponding files on disk and facilitates synchronization between the on-disk and in-memory data representations. This strategy makes them memory efficient and particularly well-suited for handling scATAC-seq data. For example, it enables ArchR and SnapATAC2 to use genome-wide bins with a size as small as 500bp. Although both Signac and SnapATAC2 demonstrate comparable memory usage based on the datasets we examined, SnapATAC2 objects tend to contain many more features than Signac objects.

Depending on the method for feature matrix construction, the running time for aggregation can vary. The aggregation steps are relatively fast, e.g. within 10 minutes for 20000 cells.

For ArchR and Signac, it depends on how the genomic features are defined. Unsurprisingly, ArchR peaks nearly doubled the running time compared to ArchR tiles, and Signac by cluster peaks doubled the time compared to Signac all cell peaks. This is because ArchR peaks/Signac by cluster peaks performed a second round of processing on top of ArchR tiles/Signac all cell peaks. ArchR peaks and Signac by cluster peaks aim to trade speed for improved identification of small cell classes.

## Discussion

We benchmarked 8 data processing pipelines derived from 5 different methods developed for scATAC-seq data, focusing on their capability to discern cell heterogeneity and delineate cell types. By using 10 metrics to assess the performance at the embedding, graph, and partition levels, we systematically examined each pipeline and evaluated the impact of key parameter choices at each data processing stage. We observed that the ranking of methods is dependent on the complexity of the datasets. For simpler datasets with distinct cell types, aggregation outperformed other methods and demonstrated superior performance in identifying small cell classes. SnapATAC2 emerged as the second-best method, while ArchR and Signac struggled to identify rare types. For complex datasets presenting hierarchical clustering structures and highly similar subtypes, SnapATAC and SnapATAC2 proved to be the most efficient in distinguishing subtypes. The aggregation method is second best, although occasionally it failed to detect differences between subtypes.

Our evaluation metrics measured the efficacy of feature engineering at each stage of the clustering pipeline, including cell embedding learning, SNN construction, and graph-based partitioning. On one hand, this approach allows us to dissect distinct facets of clustering performance. For instance, ARI2 offers a sensitive measure of the quality of rare cell type identification, while PWC, at the SNN graph level, quantifies the extent of isolation between cell types and is well-suited for evaluating both small and closely related classes. On the other hand, our metrics facilitate a rigorous evaluation that is not confounded by potentially suboptimal parameter choices at intermediate stages. For example, the AUC of ARI provides an overall performance summary across different resolutions, while cLISI assesses the cell embedding, which is at the stage prior to the determination of resolution and partitioning. The ranking of methods defined by different metrics is not always consistent with each other, which is a common observation in various benchmarking efforts. This further highlights the importance to incorporate multiple metrics and allows users to focus on the most relevant aspects of the evaluation according to their biological questions.

We built our benchmark on 6 datasets with different types of annotations serving as the ground truth. Among these, 4 datasets contain annotations from either genotype, tissue origins, or FACS labels, which we regard as high confidence for those specific datasets. The remaining 2 datasets are multi-omic data, where RNA modality is used to infer annotation. In these two datasets, we observed that the number of natural clusters does not always agree between RNA and ATAC, and the best ARI is not always achieved at the number of clusters of RNA data. While there may be discrepancies between RNA and ATAC classes, for example if epigenetic differences lack a transcriptomic correlate, these discrepancies are unlikely to significantly bias our comparisons. However, for cell state differences that are specific to epigenetic changes but lack transcriptomic alterations, our benchmark will not be able to include their comparison. In such cases, multi-omics datasets with multiple epigenomic layers might help.

A potential limitation of our study lies in the composition of our datasets, which predominantly consist of well-defined cell types rather than a spectrum of continuous cell states. This could bias our evaluation in favor of methods that facilitate clear separation between distinct states. In scenarios where mapping a continuous trajectory is critical for downstream analyses, the preferred method might differ. However, cell-type clustering remains at the heart of single-cell data analysis. Identifying the most effective feature engineering method to discern cell type differences also contributes towards establishing a foundation for studying state-informative features in future studies.

We believe that our benchmark not only provides practical guidance for users in choosing methods for their biological analyses, but also illuminates areas for potential improvements in future method development. First, while previous benchmarks have concluded that methods based on aggregating accessible chromatin regions at the motif or gene level generally underperform ^[4]^, our benchmark illustrates that a purely data-driven aggregation strategy can achieve top performance. This suggests that redundant information in scATAC-seq data exists and can be harnessed to reduce noise.

Our analysis also highlights the challenge of mitigating library size effects. Library size effects, caused by technical variations, have long been observed in next-generation sequencing data, and normalization steps to correct these are now standard practice for single-cell RNA-seq data ^[19,31,32]^. However, in the context of scATAC-seq data, this issue has not been adequately characterized or addressed. One aim of TF-IDF transformation performed in Signac is to correct for the library size difference between cells. From our observation, this is not very efficient. Linear regression-based normalization implemented in SnapATAC and SnapATAC2 seems to work well, but further comparison is needed. The binarization of peaks or bins can also be regarded as a normalization strategy ^[32]^, but recent work has also shown that retaining the count information instead of binarizing it can improve the performance of some models ^[28,33]^, indicating that even in single cells, chromatin accessibility may actually be quantitative. In summary, striking a balance between removing the technical variance and avoiding excessive correction that could mask biologically meaningful differences in global or local accessibility levels remains a challenge.

The field of computational methods for scATAC-seq data is continually advancing, with new methodologies regularly emerging. There is an ongoing need for robust and neutral benchmarking efforts that serve both method users and developers effectively. While we have incorporated the most prevalent and recent methods in this study, we acknowledge that the immediacy of our work will inevitably diminish over time. To facilitate future benchmarking work, we offer a reproducible and expandable Snakemake pipeline of our benchmarking framework, and have made our processed datasets and intermediate data publicly available.

## Conclusions

Taking together, we suggest choosing method for scATAC-seq analysis according to the complexity and size of the targeted dataset. For datasets with a simple structure where cell types are distinct from each other, all methods generally perform well; if small cell classes are expected and of interest, the aggregation method, SnapATAC, or SnapATAC2 are preferred. For more difficult tasks with hierarchical clustering structures and highly similar subtypes, SnapATAC and SnapATAC2 are among the best choice.

When the dataset is large (e.g. more than 20000 cells), SnapATAC is not very memory efficient on a typical desktop computer, and SnapATAC2 is preferred. Signac generally performs better than ArchR, but ArchR is more memory efficient. Aggregation steps do not add much time and memory consumption on top of Signac, so whenever Signac is used, the aggregation method can also be performed easily.

During the feature engineering steps, our results suggested that the choice peaks versus bins, or one-step versus two-step peak calling, are usually comparable in their performance. Users can choose according to their preferences. If SnapATAC or SnapATAC2 is used, a dimension of the latent space between 10 and 30 is recommended. For Signac and ArchR, 10 to 50 dimensions represent a reasonable range, while for aggregation, a larger number of dimensions is still suitable.

## Methods

### Datasets and preprocessing

For our benchmark, we used 6 scATAC-seq or single-cell multi-omics datasets that are publicly available ^[9,24,25,26]^ (see Table 1; links to the public repositories can be found in the “Availability of data and materials” section). For datasets where the fragment files are publicly available, these were downloaded from the author’s repositories. For datasets where the fragment files are not available, we downloaded the bam files and used the command line tool Sinto ^[34]^ to create fragment files.

For each scATAC-seq (or the ATAC component of the multi-omic) dataset, we first performed per-cell quality control (QC) using ArchR (v1.0.3) ^[9]^ by thresholding the Transcription Start Site Enrichment Score (TSSE) and the number of unique fragments; the thresholds for each dataset are in Table 2. Then we applied doublet removal procedures using `addDoubletScores()` and `filterDoublets()` in `ArchR`. Key parameters for these two functions are in Table 2, including k in function `addDoubletScores()`, and filterRatio in function `filterDoublets()`. We then filtered the fragment files to keep only cells that passed QC. For single-cell multi-omics datasets, we filtered by QC of both the ATAC and the RNA modalities. QC of RNA-seq was conducted using Seurat (v4.3.0) ^[35]^ by applying filters `nCount_RNA > 800 & percent.mt < 5` for Chen2019, and `nFeature_RNA > 200 & nFeature_RNA < 5000 & nCount_RNA < 25000 & percent.mt < 20` for 10XPBMC. Doublets were identified using function `scDblFinder()` in R package `scDblFinder` (v1.13.9) ^[27]^ and then removed. Before calling `scDblFinder()`, Louvain clustering ^[36]^ was performed with resolution= 0.5 for Chen2019 and 0.8 for 10XPBMC in Seurat. Then the clusterings results were used in `scDblFinder()`. All these filtering steps were applied to the fragment files, and the final filtered fragment files were inputs for the Snakemake pipeline.

**Table 2:** Details on scATAC-seq QC and doublet removal. NA means the corresponding step is not performed in that dataset. If a parameter of these two steps is not mentioned, then the default value was used.

| Dataset | TSSE threshold (min) | TSSE threshold (max) | nFrag threshold | k | filterRatio |
| --- | --- | --- | --- | --- | --- |
| Cell line | 1 | NA | 1000 | 50 | 3 |
| Atlas1 | 7 | NA | 1000 | NA | NA |
| Atlas2 | 7 | NA | 1000 | NA | NA |
| 10XPBMC | 8 | NA | 1000 | 30 | 1 |
| Buenrostro2018 | NA | NA | NA | NA | NA |
| Chen2019 | 2 | 20 | 500 | 30 | 1 |

### Datasets from the human adult single-cell chromatin accessibility atlas

The human adult single-cell chromatin accessibility atlas ^[24]^ contains 111 distinct cell types across 30 tissues and is a rich resource for scATAC-seq data of different cell types. We took two subsets of this atlas as our evaluation datasets “Atlas1” and “Atlas2”. The idea is to use the tissue of origin as the ground truth for benchmarking clustering. By examining the fraction of cells from different tissues for each cell type, we found that many cell types exist exclusively in one tissue (Figure S14). Therefore, we selected tissue-cell-type pairs by first selecting a subset of cell types that have *≥* 85% of cells from the same tissue, and then for each of the corresponding tissues, we selected one cell type randomly, but excluded cell types that have less than 300 cells. For each tissue with multiple samples, we selected one sample that contains the maximum number of cells of that cell type that have passed QC. We used only one sample for each cell type to eliminate any potential batch effect. For the tissue-cell-type pairs selected for each dataset, we downsampled cells per cell type, using fractions 0.3 for Atlas1 and 0.5 for Atlas2. The tissue-cell-type pairs and the sampled cell ID for “Atlas1” and “Atlas2” are available on GitHub ^[37]^.

### scRNA-seq data annotation

For 10XPBMC and Chen2019 datasets, the RNA modality was subjected to Leiden clustering in Seurat using resolution=0.5 for Chen2019 and resolution=0.8 for 10XPBMC. Then for each cell, a label was transferred by reference mapping ^[35]^ using Seurat. In the case of the 10XPBMC dataset, an annotated PBMC reference dataset ^[38]^ was utilized for label transfer. As for the Chen2019 dataset, the scRNA-seq data of the adult mouse brain from the Allen Brain Atlas ^[39]^ served as the reference dataset. Then we performed some manual curation to get the final cluster annotation. We describe roughly this process below. For each Leiden cluster, the majority cell label was token as the label of that cluster. If two clusters got the same label, we subset all cells from these two clusters and performed multiple rounds of clusterings on this subset. If the splitting of these two clusters were stable, we labeled them differently, including “CD4 Naive 1” and “CD4 Naive 2”. Otherwise, we merged these two clusters.

### Feature engineering methods

For all methods, we followed the procedures recommended in the author’s documentation.

### Signac

Starting from the fragment file, Signac first uses MACS2 for peak calling, then performs LSI on the peak count matrix to obtain a low-dimensional representation. Peak calling was conducted in two ways: (1) aggregating all cells for peak calling (denoted as “Signac all cell peaks”), or (2) aggregating cells and calling peaks for each cluster individually, followed by generating a consensus peak set from the peaks identified in all clusters, referred to as “Signac by cluster peaks.” LSI consists of 3 steps: (1) normalization using term frequency-inverse document frequency (TF-IDF); (2) selecting the top 95% most common peaks; (3) performing singular value decomposition (SVD).

We used the R package Signac (v1.9.0) for its implementation. As suggested by the tutorial: https://stuartlab.org/signac/articles/pbmc_multiomic.html, we created a fragment object, called MACS2, and removed peaks on nonstandard chromosomes and genomic blacklist regions, and peaks having width *<* 20 or width *>* 10000. To identify peaks per cluster, the cell embeddings generated by “Signac all cell peaks” are used to define clusters using the Louvain algorithm with a default resolution of 0.8.

Subsequently, we performed normalization, feature selection, and linear dimensional reduction using `RunTFIDF()` with `method=1, FindTopFeatures()` with `min.cutoff=“q5”`, and `RunSVD()` with n=100, respectively. As suggested in the tutorial, the first LSI components often capture sequencing depth. We removed LSI components that have larger than 0.75 Pearson correlation with the total number of counts.

### Aggregation

The aggregation method starts with the peak count matrix where the peak set is identified using the method “Signac by cluster peaks”. Then the cell-by-peak fragment count matrix is used for subsequent TF-IDF normalization and PCA. Minibatch K-means clustering is applied to the PCA to cluster peaks into meta-features (K=1000 by default). Ultimately, an aggregated count matrix is obtained by summing the counts per meta-feature, and PCA is performed on the aggregated count matrix to get the low-dimensional representation.

We used the function `aggregateFeatures()` in `R` package scDblFinder (v1.13.9) with the default parameters. By default, *K* = 1000 feature clusters are identified.

### ArchR

ArchR takes the fragment files as input, and can use either the genomic tiles or peaks as features. We implemented both options. The “ArchR tiles” method uses 500bp non-overlapping genomic tiles to construct a binarized tile matrix, and then performs iterative LSI on the matrix to extract meaningful low-dimensional representations. Similar to “Signac by cluster peaks” approach, “ArchR peaks” method first uses the latent representation obtained from “ArchR tiles” for clustering, then performs peak calling per individual clusters and generates a consensus peak set by merging these peak tracks. Afterwards, iterative LSI is performed on the peak count matric.

During the iterative LSI process, at each iteration, the top accessible features (in 1st iteration) or top variable features (since 2nd iteration) are selected for LSI. The resulting cell clusters are then identified and utilized for feature selection in the subsequent iteration, enabling an iterative refinement of the LSI procedure.

We used the `R` package ArchR(v1.0.3) for implementation. When running the function `addIterativeLSI()`, Louvain algorithm was used for the clustering in intermediate steps with increasing resolutions, and no subsampling of cells was performed. Other parameters we used were set to be the default values.

### SnapATAC

The SnapATAC method (version 1) takes the fragment files as input, and first constructs a binary cell-by-bin matrix using 5000bp non-overlapping genomic bins. Then after filtering out unwanted bins, it computes a pairwise cell-to-cell similarity matrix using Jaccard coefficient. This kernel matrix is subject to normalization of the coverage bias and then eigenvalue decomposition (EVD) to get the cell embeddings.

For the implementation, we used the command line tool snaptools (v1.4.8) to create snap files from fragment files, and the R package SnapATAC (v1.0.0) for the rest of the processing pipeline. We followed the standard procedures in https://github.com/r3fang/SnapATAC/tree/master/examples/10X_PBMC_15K, except that we ran the function `runDiffusionMaps()` using all cells instead of using landmark cells.

### SnapATAC2

SnapATAC2 is the version 2 of SnapATAC method and it is released as a python package. By implementing AnnData object and optimizing the on-disk representation, it facilitates the processing of high-dimensional data. As demonstrated in the tutorial https://kzhang.org/SnapATAC2/tutorials/pbmc.html, SnapATAC2 first creates a cell-by-bin matrix containing insertion counts using 500-bp bins by default. Then a pairwise cell-to-cell similarity matrix is generated, using either Jaccard coefficient (SnapATAC2 jaccard) or cosine similarity (SnapATAC2 cosine). With this kernel matrix, the symmetric normalized graph Laplacian is computed, and the bottom eigenvectors of the graph Laplacian is used as the lower dimensional representation. For implementation we used SnapATAC2 (v2.2.0). To select features, we removed bins overlapping with the blacklist regions as always done in other methods, and called function `snapatac2.pp.select_features()` with parameters `min_cells=10, most_variable=1000000`.

### Clustering

In this study, we used a well-established graph-based clustering method for all clustering analyses. We first constructed a shared nearest neighbor graph, and then applied the Leiden algorithm ^[20]^ using modularity as the optimization objective, as implemented in the Seurat package (v4.3.0) ^[35]^. The Leiden algorithm incorporates a step where node partitions are refined by randomly reassigning nodes to communities that increase the objective function, enabling a wider exploration of the partition space ^[20]^. To account for the inherent stochasticity in the Leiden algorithm, we ran it with 5 different random seeds: 0, 2, 5, 42, and 123. Since the optimal number of clusters is not known *a priori*, a range of resolutions was used to obtain diverse clustering solutions yielding varying numbers of clusters.

### Evaluation metrics

According to the data structure our evaluation applied to, we have classified our evaluation metrics into three categories: embedding-based, graph-based and partition-based (Figure 1).

### ASW, FNS

The silhouette width quantifies the average distance between an observation and the other observations within its cluster, relative to the average distance to the nearest neighboring cluster ^[40]^. The Average Silhouette Width (ASW) is calculated as the mean silhouette width across all observations within a cluster, providing insights into the compactness of the cluster and its separation from other clusters. ASW values range from -1 to 1, with 1 indicating dense and well-separated clusters, 0 representing clusters that overlap, and -1 indicating significant misclassification, where within-cluster dispersion is greater than between-cluster dispersion.

A limitation of ASW is that it is not invariant to the scaling of the space. As a solution, we introduced the fraction of negative Silhouette score (FNS) to assess the cluster-level proportion of cells with a negative Silhouette width. FNS characterizes the fraction of cells with a smaller distance to cells within another cluster compared to their own cluster. It is robust to linear scaling and enables more meaningful comparisons across different dimensional reduction methods.

### cLISI

The LISI has been proposed to evaluate either the mixing between batches or the separation between cell types ^[41]^. To calculate it, a weighted k-nearst neighbor (kNN) graph is first generated based on Euclidean distance within an embedding space. Subsequently, for each node in the graph, it computes the expected number of cells needed to be sampled before two cells are drawn from the same batch/clusters within its neighborhood. We used the cluster-based variant of LISI, known as cluster LISI (cLISI), as a metric to assess the embedding representation. cLISI ranges from 1 to K, where K is the total number of cell types in the dataset. 1 indicates a neighborhood consisting exclusively of cells from a single cell type, while N corresponds to complete mixing, with cells from all cell types found within the neighborhood.

### PWC

Partition-based metrics are susceptible to the influence of clustering parameters, whereas embedding-based metrics rely on the proper definition of similarity within the embedding space, which may not necessarily align with the similarity employed in clustering. Therefore, we proposed a novel graph-based metric that directly operates on the graph where cells of the same (ground truth) type are identified as communities. Filippo *et al*. ^[42]^ discussed a definition of community in the network by splitting the total degree of a node *i* into two contributions: given a subgraph *V* ⊂ *G*, 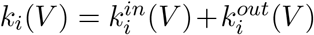, where 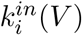 is the number of edges connecting node *i* to other nodes in *V*, and 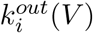 is the number of connections towards the rest of the network. The subgraph *V* is a community in a strong sense if 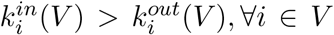. Inspired by this definition, we introduced the metric Proportion of Weakly Connected cells (PWC). PWC quantifies, for a subgraph *V* consisting of all the cells of the same true class, the proportion of cells that have fewer connections within *V* than with the rest of the graph.

### AW, AV

Wallace ^[43]^ proposed two asymmetric indices to quantify the similarity between two partitions of a set. Let *U* ={*U*_1_, *U*_2_, …, *U*_*I*_*}* be the partition of the dataset defined by cell types, and *Z* ={*Z*_1_, *Z*_2_, …, *Z*_*J*_ *}* be the partition given by the clustering prediction. The first index W is the proportion of joint object pairs in partition *U* that are also joined in partition *Z*. The second index V is the proportion of joint object pairs in partition *Z* that are also joined in partition *U*. Both index W and V can be adjusted for chance using formula:

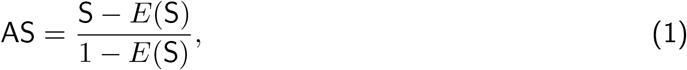

where S is a similarity measure that does not have value 0 under statistical independence. A generalized hypergeometric model is assumed to calculate the expectation value of V and W ^[44]^. AW can be interpreted as the completeness of cell types. It quantifies to what extend objects belonging to the same cell type in *U* are assigned to the same cluster in Z. Similarly, AV can be interpreted as the homogeneity of clusters, which measures to what extend clusters are not mixing objects of different cell types. AW and AV can be decomposed into indices for the individual cell types of partitions *U* and for the individual clusters of partitions Z, respectively ^[45]^, that is:

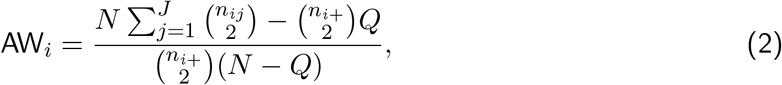

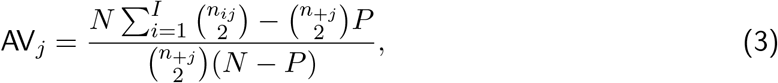

where *n*_*ij*_ is the number of objects placed in class *U*_*i*_ and in cluster *Z*_*j*_, *N* is the total number of pairs of objects, 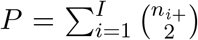 is the number of object pairs that were placed in the same cluster in *U*, and 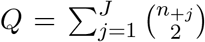 is the number of object pairs that were placed in the same cluster in *Z*. AW and AV range from *−*1 to 1, with 0 for random assignments, and 1 for perfect agreement.

### ARI, ARI2

The adjusted Rand index is the harmonic mean of AW and AV, and therefore a summary of both the homogeneity of predicted clusters and the completeness of true classes. ARI can be decomposed into a weighted average of the AW_*i*_’s and AV_*j*_’s as follows ^[45]^:

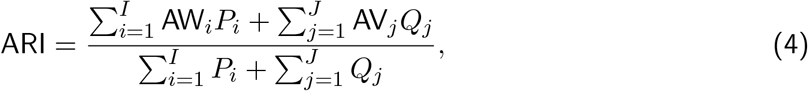

where 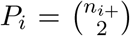 is the number of object pairs in cluster *U*_*i*_, and 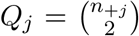 is the number of object pairs in cluster *Z*_*j*_. Equation 4 shows that ARI is largely determined by the AW_*i*_ and AV_*j*_ values of large clusters. However, in many cases in single-cell analysis, the rare cell types are of more concern. Therefore, we included a variant of ARI (ARI2) proposed by Matthijs *et al*. ^[45]^ to alleviate the class size bias of ARI.

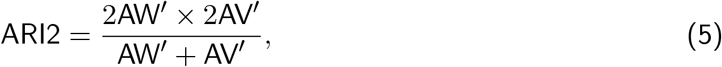

where

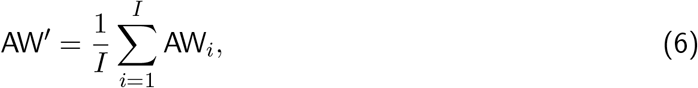

and

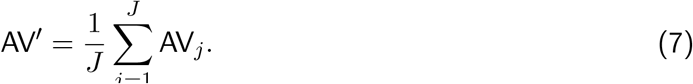

### MI, VI

While ARI, AW and AV are external evaluation metrics that count pairs of objects, mutual information (MI) and variation of information (VI) are based on information theory. These two groups of metrics do not always show consistent results, due to different underlying assumption. The MI between two partitions *U* and *Z* is as follows:

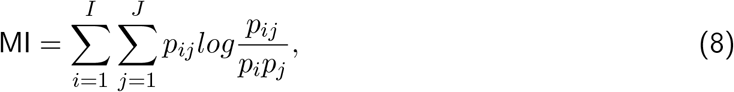

where 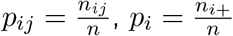, and 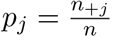. It has been shown ^[46]^ that

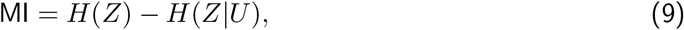

where *H*(*·*) is the Shannon entropy. Since *Z* stays the same for a given dataset, equation 9 indicates that comparing MI between methods on the same dataset is equivalent to comparing the conditional Shannon entropy of *Z* on *U*. In other words, MI can be interpreted as the measure of homogeneity of clusters, similar to AV. Note that MI is not normalized and the upper bound varies across datasets. It is therefore only meaningful to compare MI within the same dataset.

VI measures the amount of information that is lost or gained in changing from partition *Z* to *U* :

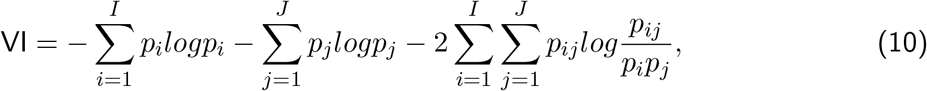

Similarly, VI is also highly related to entropy:

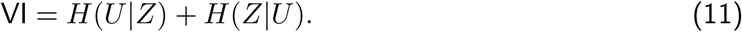

VI is not normalized, and a higher VI value indicates a worse clustering solution.

### Calculating and comparing the area under the curve

Partition-based metrics change as the clustering resolution changes. The true cluster numbers is not predetermined, and the optimal performance is not always achieved at the true number of clusters. Therefore, comparing clusterings at a fixed resolution or number of clusters becomes challenging. To address this challenge, we compared clusterings across a range of resolution parameters that result in varying number of clusters (Figure S15a). We examined the performance as the number of clusters changes and summarized the results using the area under the curve (AUC).

To calculate the AUC and compare between results of different ranges of cluster numbers, the upper bound of each metric is used for the normalization of the absolute AUC. Specifically, metrics such as ARI, ARI2, AW, and AV have an upper bound of 1. MI and VI were normalized using the empirical maximum value per method per dataset. Notably, in the case of VI, instead of using the normalized AUC directly for comparison, we used 1*−* normalized AUC.

When plotting the AUC heatmap, we colored the heatmap using the deviations from the column-wise median scaled by matrix-wise median absolute deviation. Let ***A*** be the original matrix storing the metric values, ***B*** be the transformed matrix, and *A*_*i,j*_, *B*_*i,j*_ is the element of matrix ***A***, and ***B***, respectively. The calculation of ***B*** is as in Equation 12, 13 and 14. By applying this transformation, the color scale is unified across datasets.

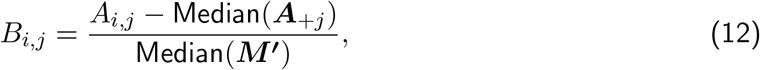

where

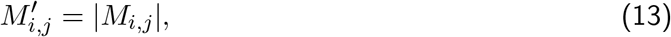

and

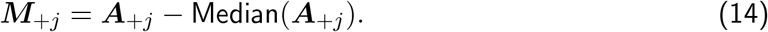

### Choosing the number of dimensions

When applying dimensional reduction methods, a parameter that one needs to choose is the number of dimensions *n* of the embedding space to use. Since all the methods use either principle component analysis (PCA) or singular value decomposition (SVD), we applied the elbow approach and examined the scree plot of each method by plotting the proportion of variance explained by each component against the component indices. We observed that for nearly all methods and datasets, the elbow point is before 15 dimensions (Figure S15b). We therefore used *n* = 15 for all methods. More details on how *n* affects the performance is discussed in Results.

## Other indices

### Evenness

Evenness (E) quantifies the homogeneity of abundances of different types in a sample ^[47]^. Here we use Equation 15 to calculate E:

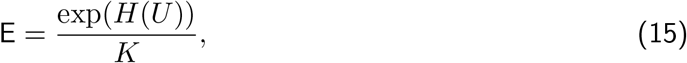

where *H*(*·*) is Shannon entropy, and *K* is the total number of cell types. E ranges from 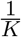 to 1, and a higher E indicates that the dataset is more balanced.

### Geary’s C

We calculated Geary’s C index ^[48]^ of log-transformed fragment counts using spatial distance defined by k-nearest neighbor (KNN) graph (k=20) ^[14]^. Geary’s C is calculated as:

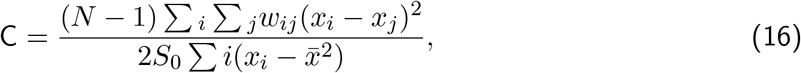

where N is the total number of cells; *x*_*i*_ is the log-transformed fragment counts of cell *i, w*_*i*_*j* is the weight of edge between cell *i* and *j* on the KNN graph, and *S*_0_ is the sum of all weights in ***W***.

## Acknowledgements

We thank all members of the von Meyenn group and Robinson group for helpful discussions and support. We specifically thank Adhideb Ghosh for advice on the study design and scRNA-seq data analysis, João Pedro Agostinho de Sousa for help with setting up the computational environments, and Emanuel Sonder for his feedbacks on the manuscript draft.

## Availability of data and materials

Our benchmarking workflow is provided as a reproducible Snakemake pipeline on GitHub: https://github.com/RoseYuan/sc_chromatin_benchmark ^[49]^. Notebooks, R scripts and supporting data used for preprocessing datasets and generating all the visualizations in this manuscript is available at https://github.com/RoseYuan/benchmark_paper^[37]^. For the analyzed datasets, the preprocessed data that can be directly input into the Snakemake pipeline is available on Zenodo ^[50]^. For the unprocessed data, fragment files of the cell line dataset were downloaded from GEO accession GSE162690, fragment file and the gene expression matrix file of the 10X PBMC multiomics dataset were downloaded from https://www.10xgenomics.com/resources/datasets/pbmc-from-a-healthy-donor-granulocytes-removed-through-cell-sorting-10-k-1-standard-1-0-0, and fragment files of the human adult atlas datasets were downloaded from GEO accession GSE184462. Bam files of the dataset Buenrostro2018 was downloaded from GEO accession GSE96772. Bam file of the ATAC part of Chen2019 dataset was processed by Stuart’s lab and we downloaded the fragment files they provided at https://stuartlab.org/signac/articles/snareseq.html, and the RNA part was downloaded from GEO accession GSE126074. The Seurat objects of the Allen mouse brain reference dataset used for annotating the scRNA-seq data were downloaded from Signac’s website: https://signac-objects.s3.amazonaws.com/allen_brain.rds. The PBMC reference dataset was downloaded here: https://atlas.fredhutch.org/data/nygc/multimodal/pbmc_multimodal.h5seurat.

## Authors’ contributions

SL and FvM conceptualized the study with the help from MDR and PLG. SL designed the benchmark, prepared the data, wrote the code for the pipeline, and performed the analysis. SL, PLG, MDR and FvM interpreted the results. SL wrote the manuscript draft. All authors reviewed and approved the final version of this manuscript.

## Authors’ information

Laboratory of Nutrition and Metabolic Epigenetics, Institute for Food, Nutrition and Health, Department of Health Sciences and Technology, ETH Zurich, Zurich, Switzerland Siyuan Luo & Ferdinand von Meyenn

Department of Molecular Life Sciences, University of Zürich, Zürich, Switzerland Siyuan Luo, Pierre-Luc Germain & Mark D. Robinson

SIB Swiss Institute of Bioinformatics, Zürich, Switzerland Pierre-Luc Germain & Mark D. Robinson

D-HEST Institute for Neurosciences, ETH Zürich, Zürich, Switzerland Pierre-Luc Germain

## Supplementary Figures

**Figure S1:**
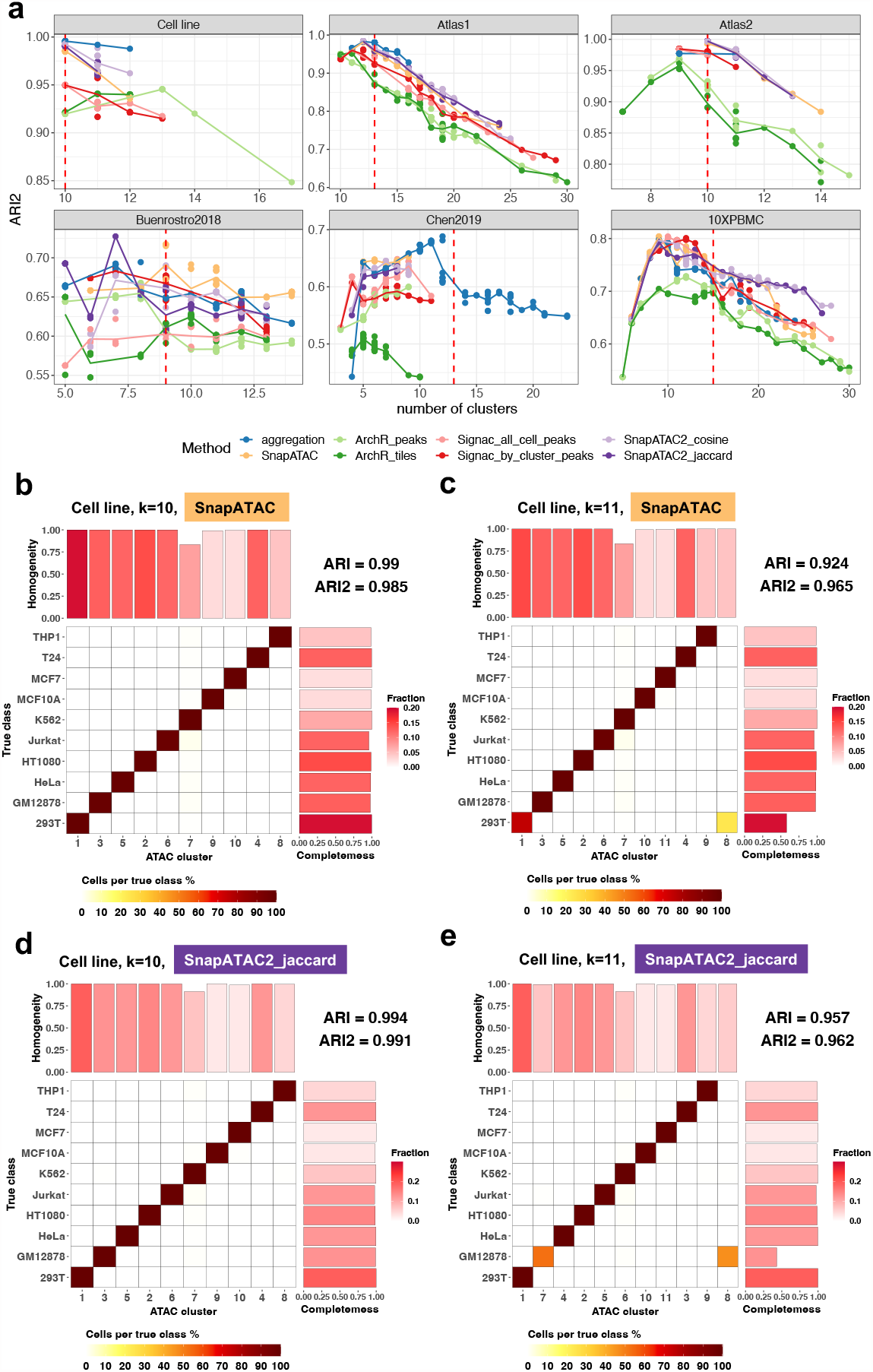
**a** ARI2 plotted against various number of clusters; as shown in Figure 3a, each subpanel represents a dataset. Each point represents a clustering solution obtained by varying the resolution parameter and the random seed in Leiden algorithm. The line plot is the average ARI2 at a given number of clusters. **b-e** True classes and their fractions of agreement with the predicted clusters. **b-c** are SnapATAC on dataset Cell line, and **d-e** are SnapATAC2 jaccard on Cell line. The x-axis is the predicted clusters, and the y-axis is the ground truth classes. The colors of tiles indicate the proportion of cells from the corresponding true class (each row sums up to one). A clearer diagonal structure indicates better agreement. ARI and ARI2 are calculated and shown on the top right. The barplot on top shows the value of AV (Methods) and can be interpreted as the homogeneity of the corresponding clusters. The barplot on the right shows the value of AW and represents the completeness of each true class in the prediction. The color of the bars shows the proportion of cells in each cluster/ground truth class. In title, the corresponding datasets, methods, and number of clusters are indicated.

**Figure S2:**
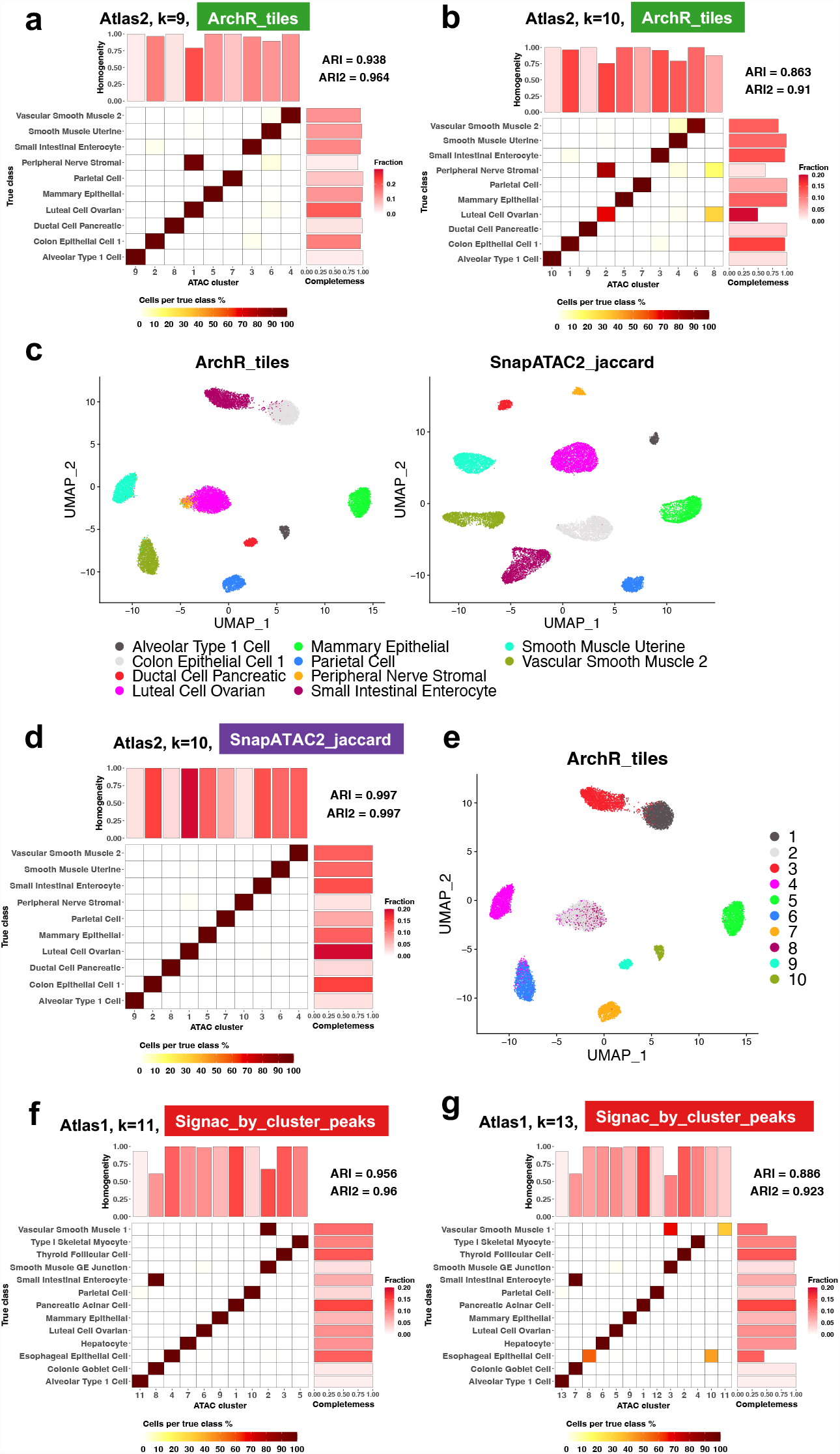
**a,b** Clustering solutions by ArchR tiles on Atlas2, with different numbers of clusters identified. **c** UMAP of Atlas2 generated by ArchR tiles and SnapATAC2 jaccard, annotated by the true classes. **d** The corresponding clustering solutions by SnapATAC2 jaccard on Atlas2. **e** UMAP annotated by the clustering solutions of ArchR tiles in **b**. Cluster 8 is the wrongly identified clusters. **f,g** Clustering solutions by Signac by cluster peaks in dataset Atlas1, with different numbers of clusters identified.

**Figure S3:**
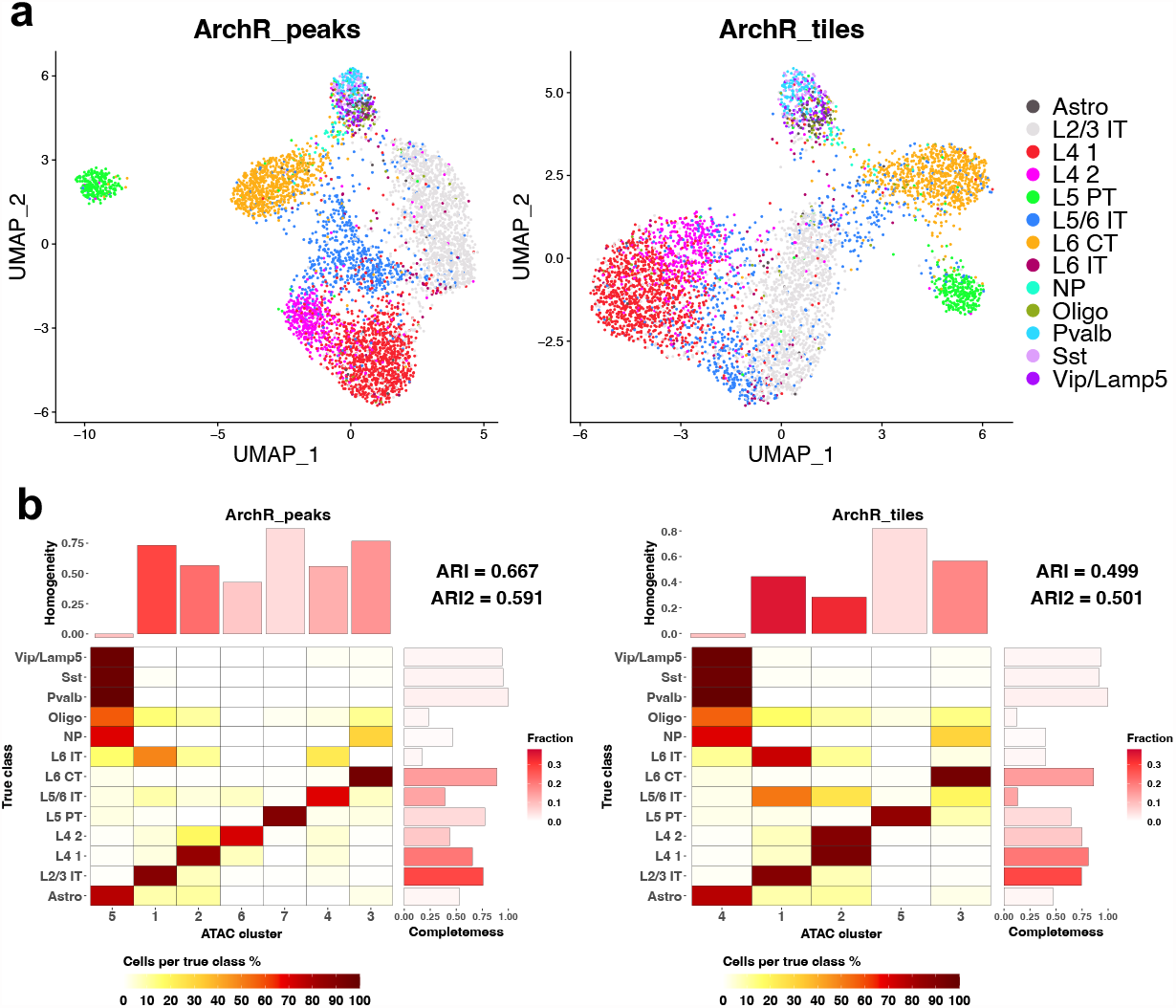
**a**UMAP of Chen2019 generated by ArchR peaks and ArchR tiles. **b** Clustering solutions of the highest ARI by ArchR peaks and ArchR tiles on Chen2019.

**Figure S4:**
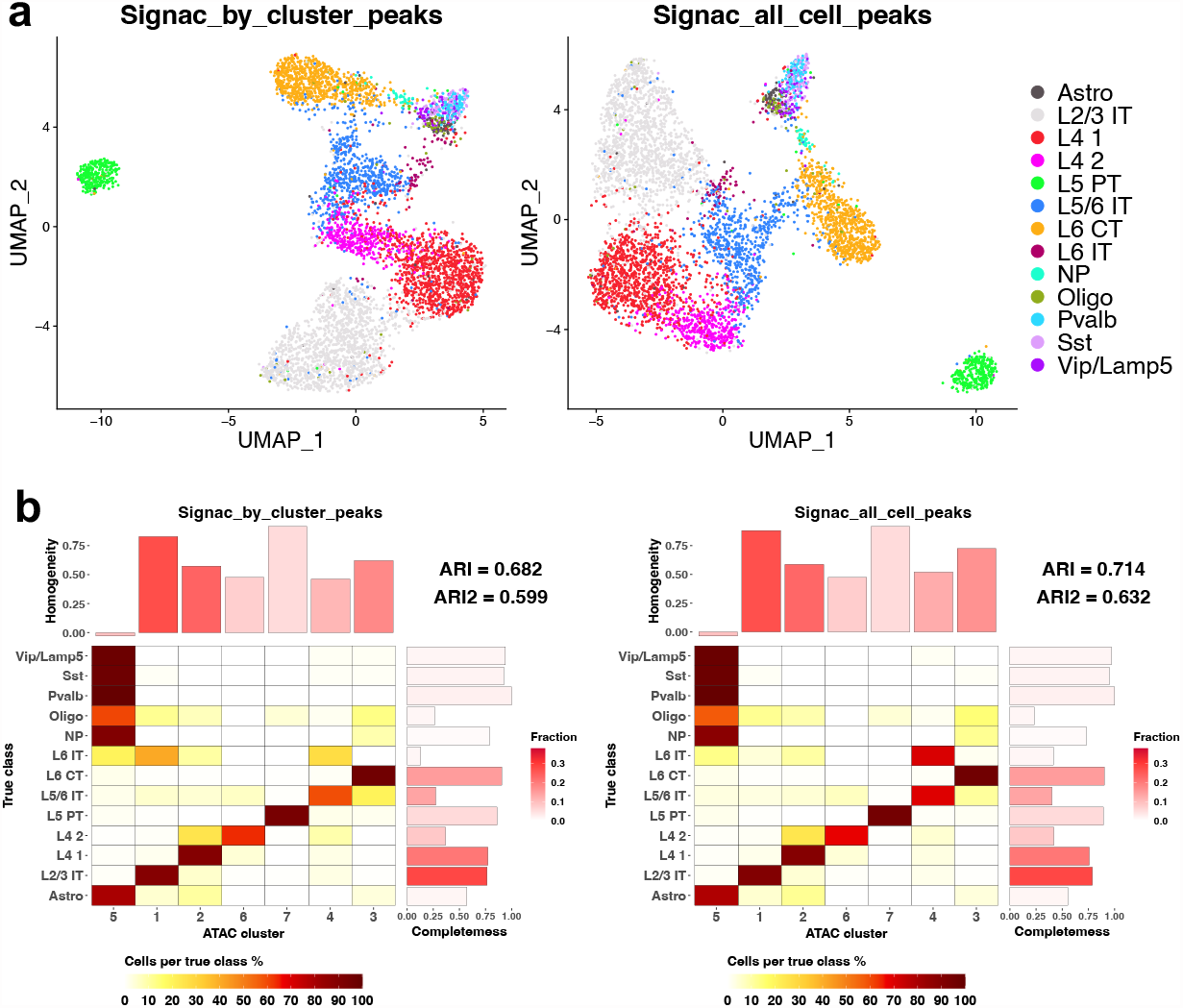
**a**UMAP of Chen2019 generated by Signac by cluster peaks and Signac all cell peaks. **b** Clustering solutions of the highest ARI by Signac by cluster peaks and Signac all cell peaks on Chen2019.

**Figure S5:**
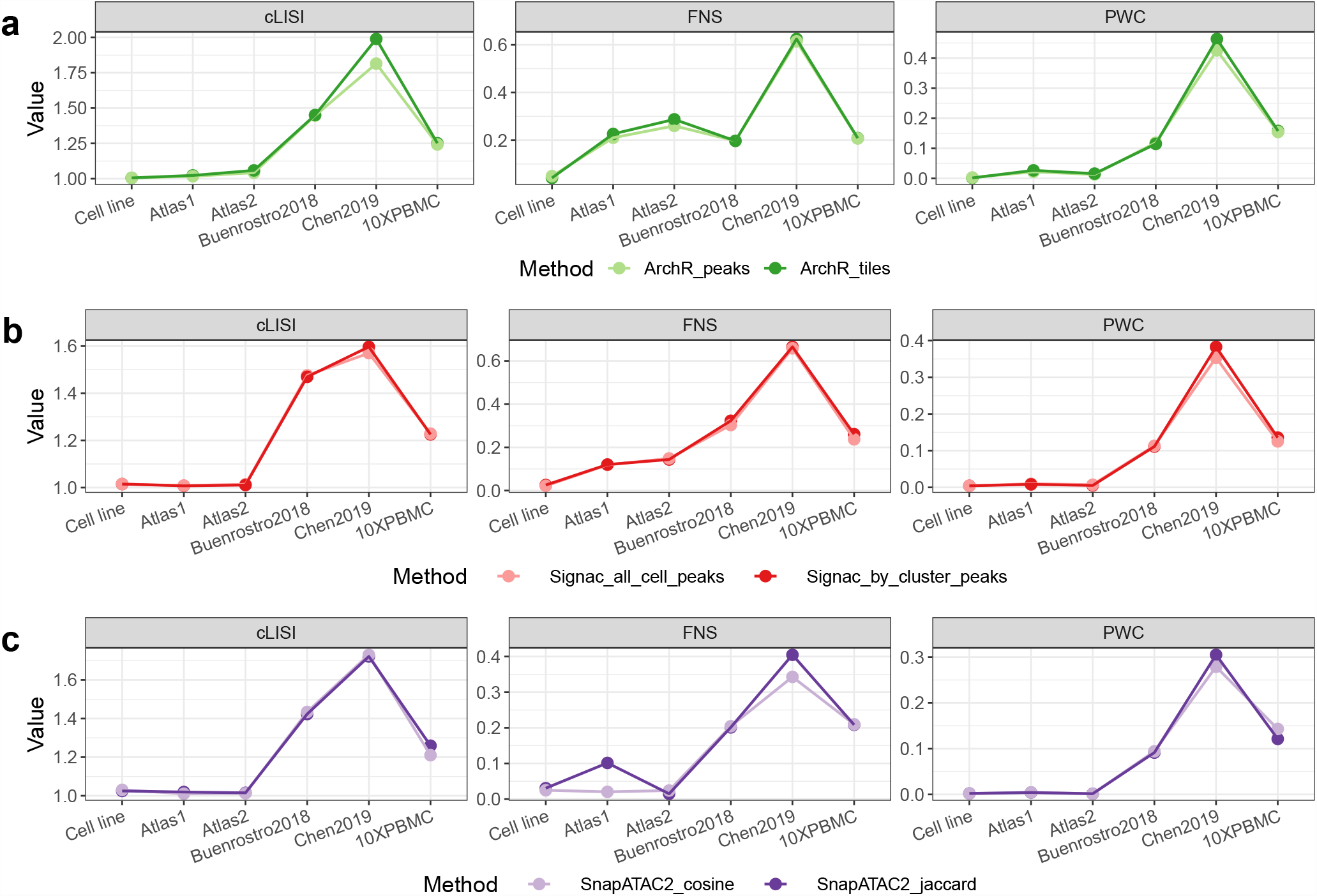
Comparison of the method performance between different data processing choices, measured by the embedding and graph-level metrics. **a** Comparing ArchR peaks and ArchR tiles. **b** Comparing Signac all cell peaks and Signac by cluster peaks. **c** Comparing SnapATAC2 cosine and SnapATAC2 jaccard.

**Figure S6:**
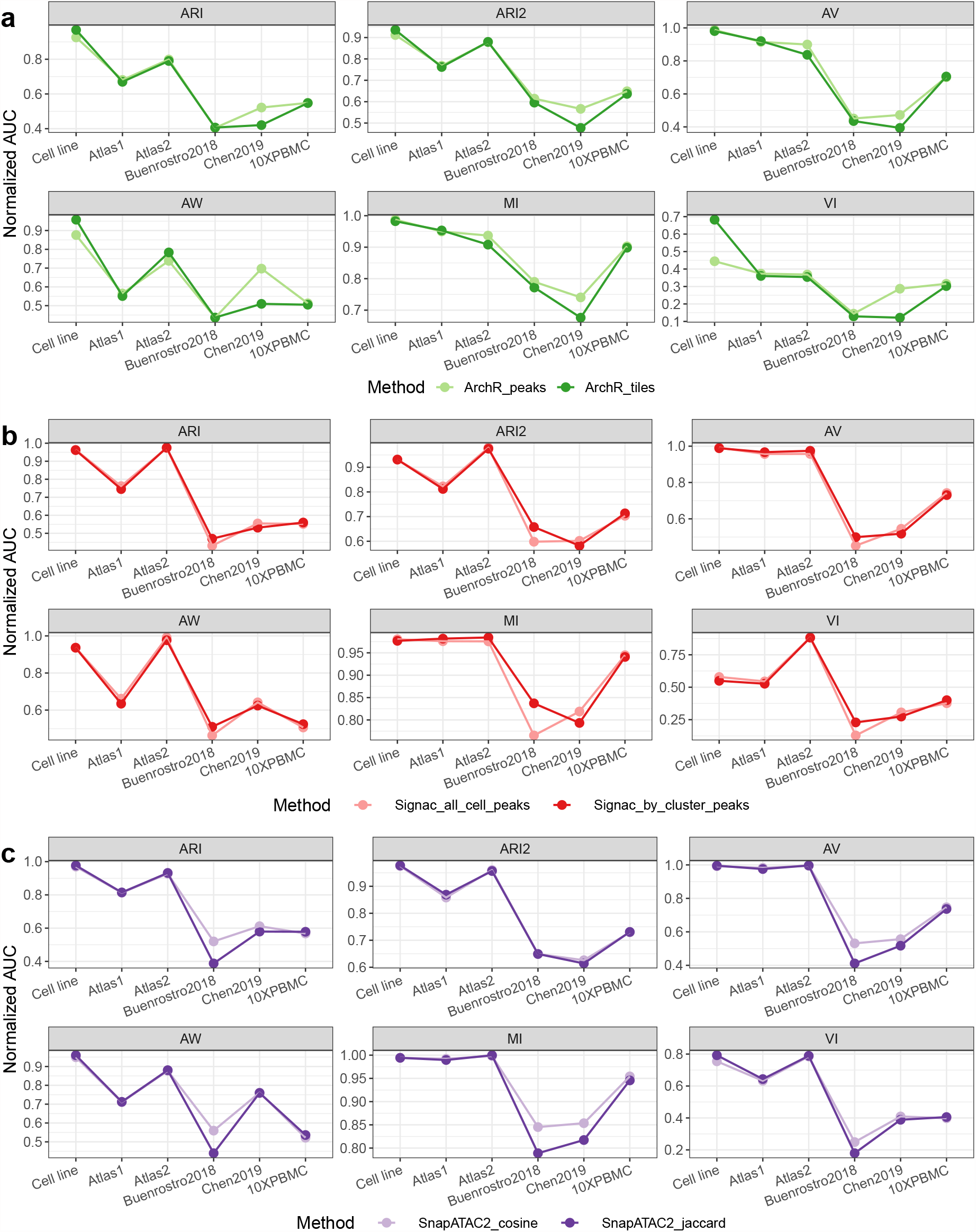
Comparison of the method performance between different data processing choices, measured by the partition-level metrics. **a** Comparing ArchR peaks and ArchR tiles. **b** Comparing Signac all cell peaks and Signac by cluster peaks. **c** Comparing SnapATAC2 cosine and Snap-ATAC2 jaccard.

**Figure S7:**
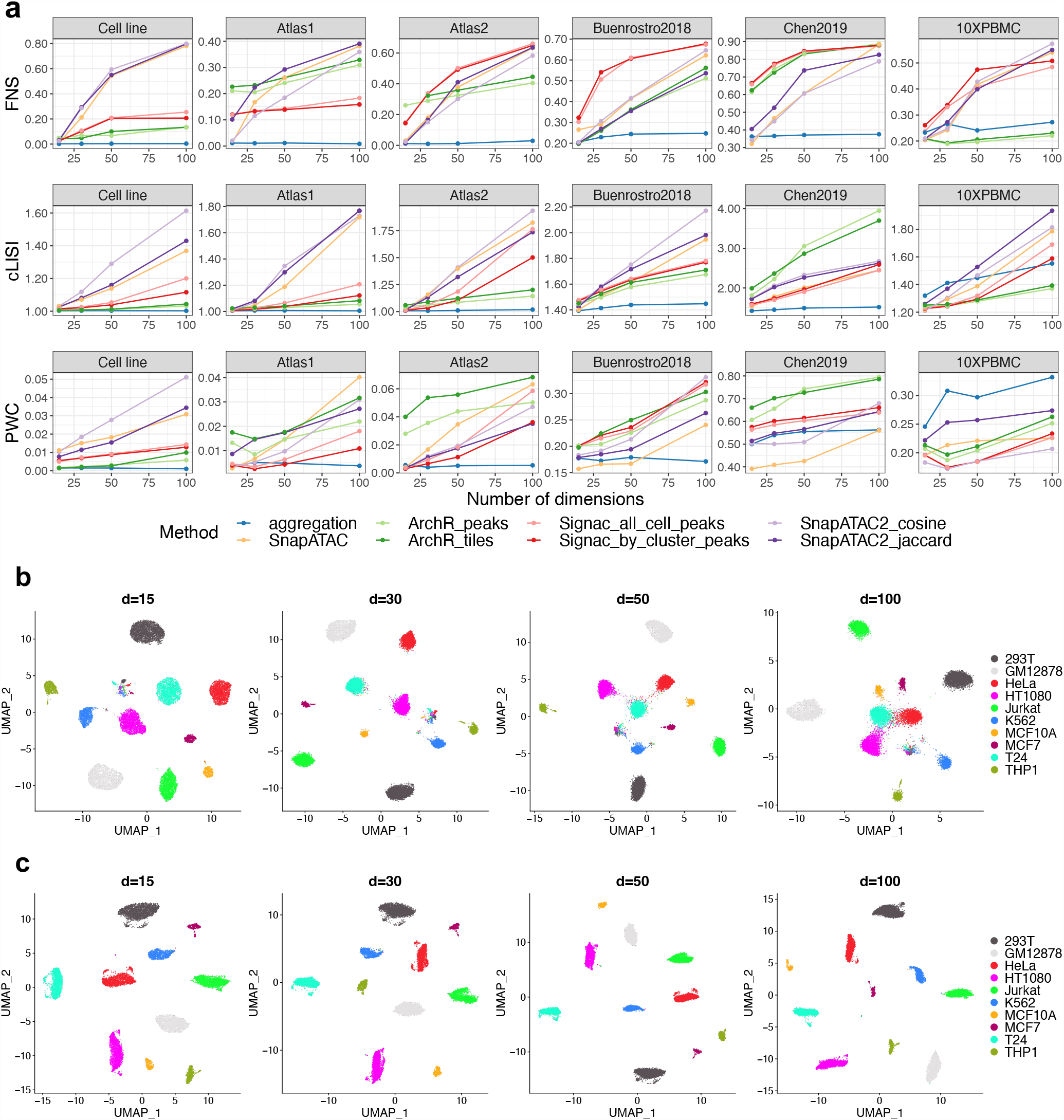
**a** Performance of each method across different choices of the number of latent dimensions, evaluated using embedding-level and graph-level metrics. **b** UMAP of Cell line dataset using SnapATAC2 cosine method. **c** UMAP of Cell line dataset using aggregation method.

**Figure S8:**
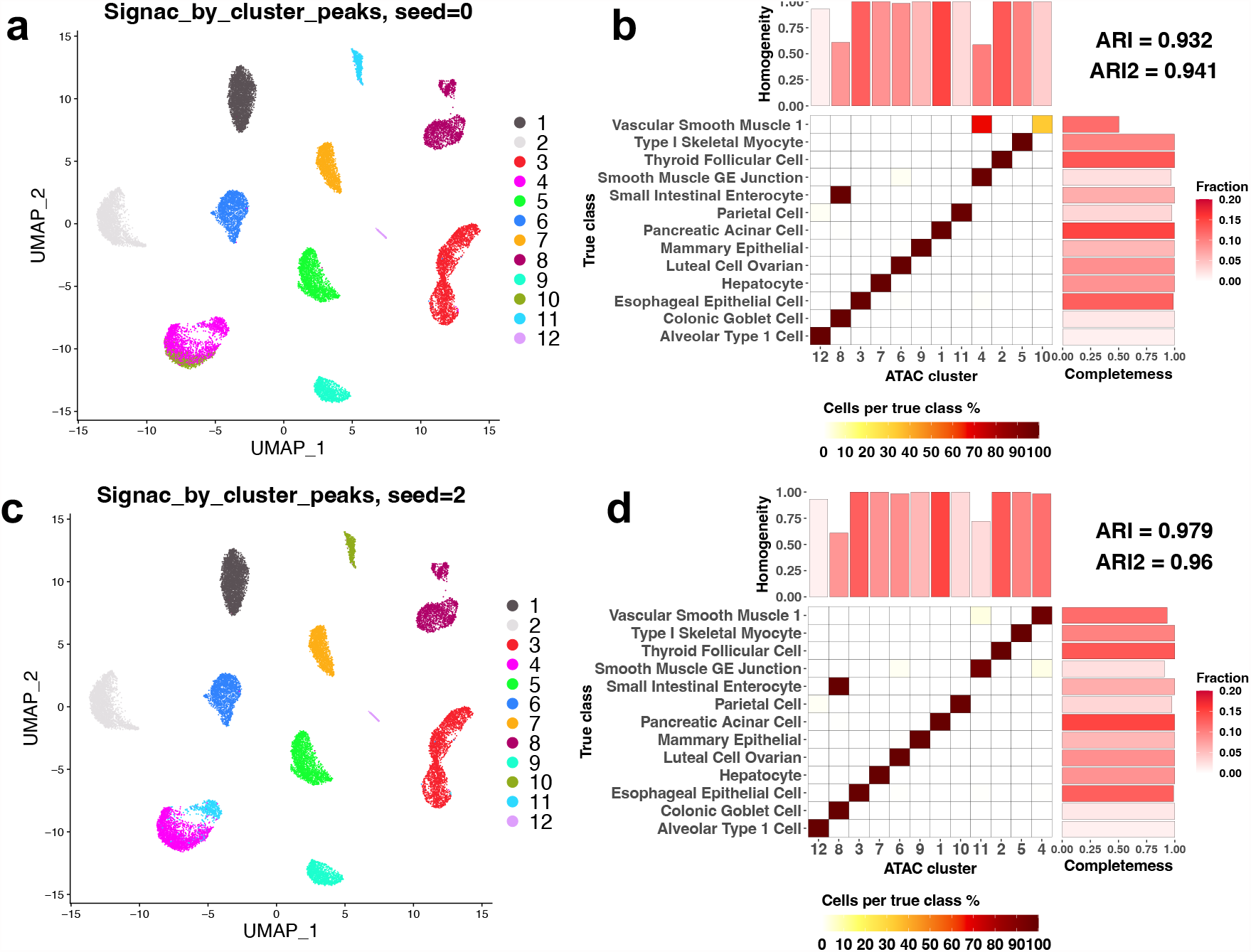
Clustering solutions of Signac_by_cluster_peaks on Atlas1, using random seed 0 (**a**) or random seed 2 (**b**) when running Leiden algorithm.

**Figure S9:**
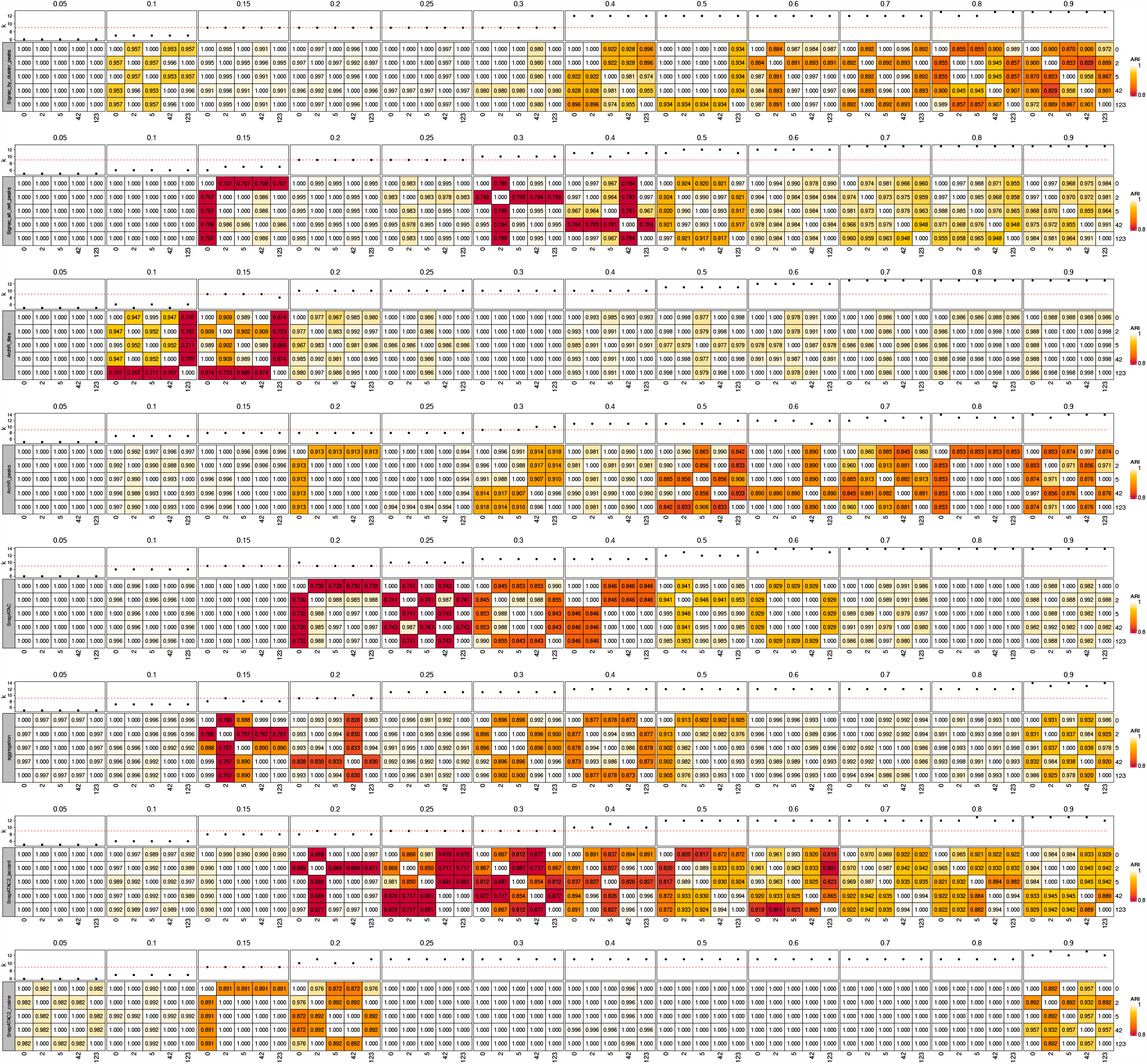
ARI between clustering solutions using the 5 random seeds, in combination with different methods and resolution parameters in dataset Buenrostro2018. The dot plots on top show the number of clusters, and the red horizontal lines are the ground truth cluster numbers.

**Figure S10:**
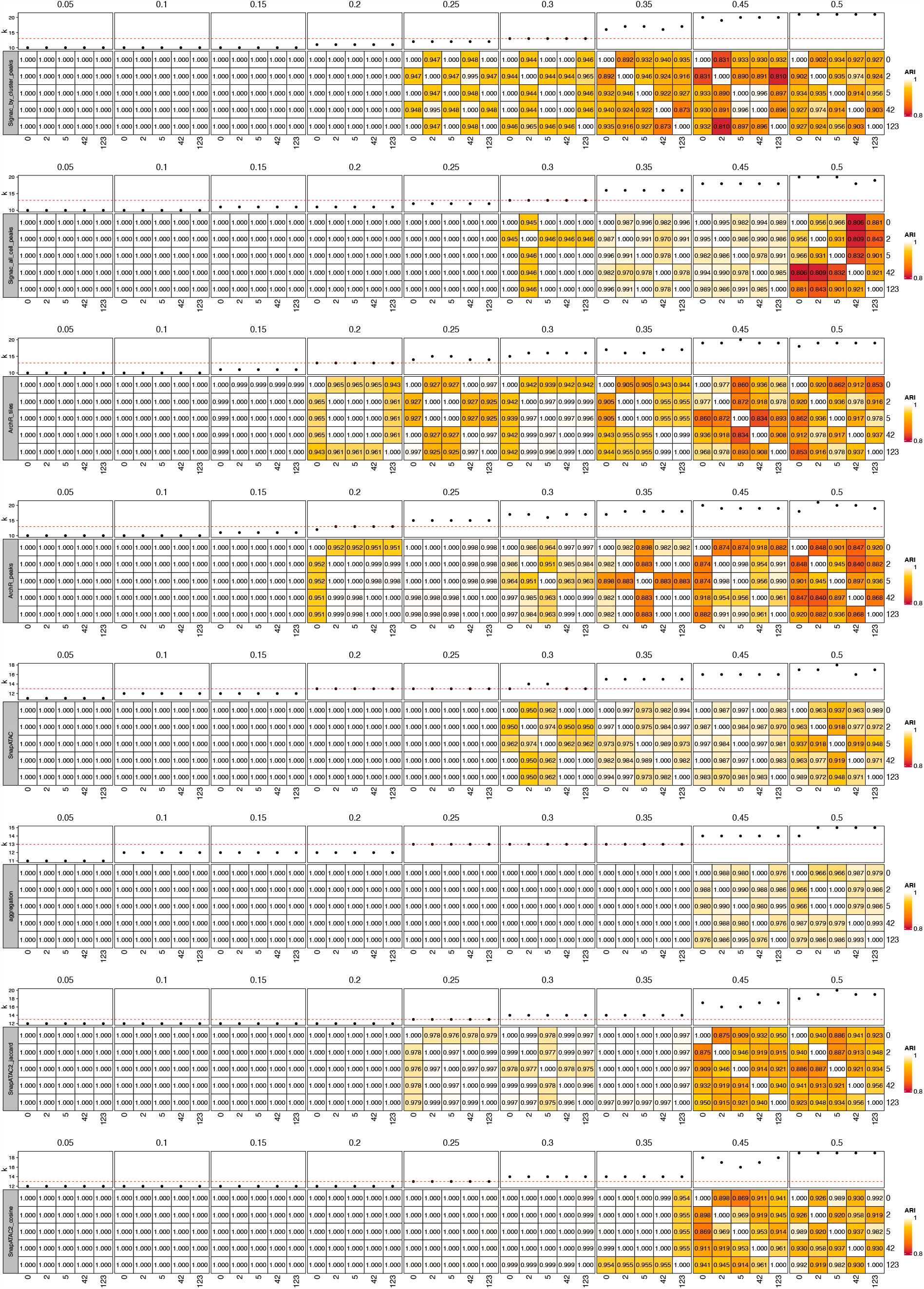
ARI between clustering solutions using the 5 random seeds, in combination with different methods and resolution parameters in dataset Atlas1. The dot plots on top show the number of clusters, and the red horizontal lines are the ground truth cluster numbers.

**Figure S11:**
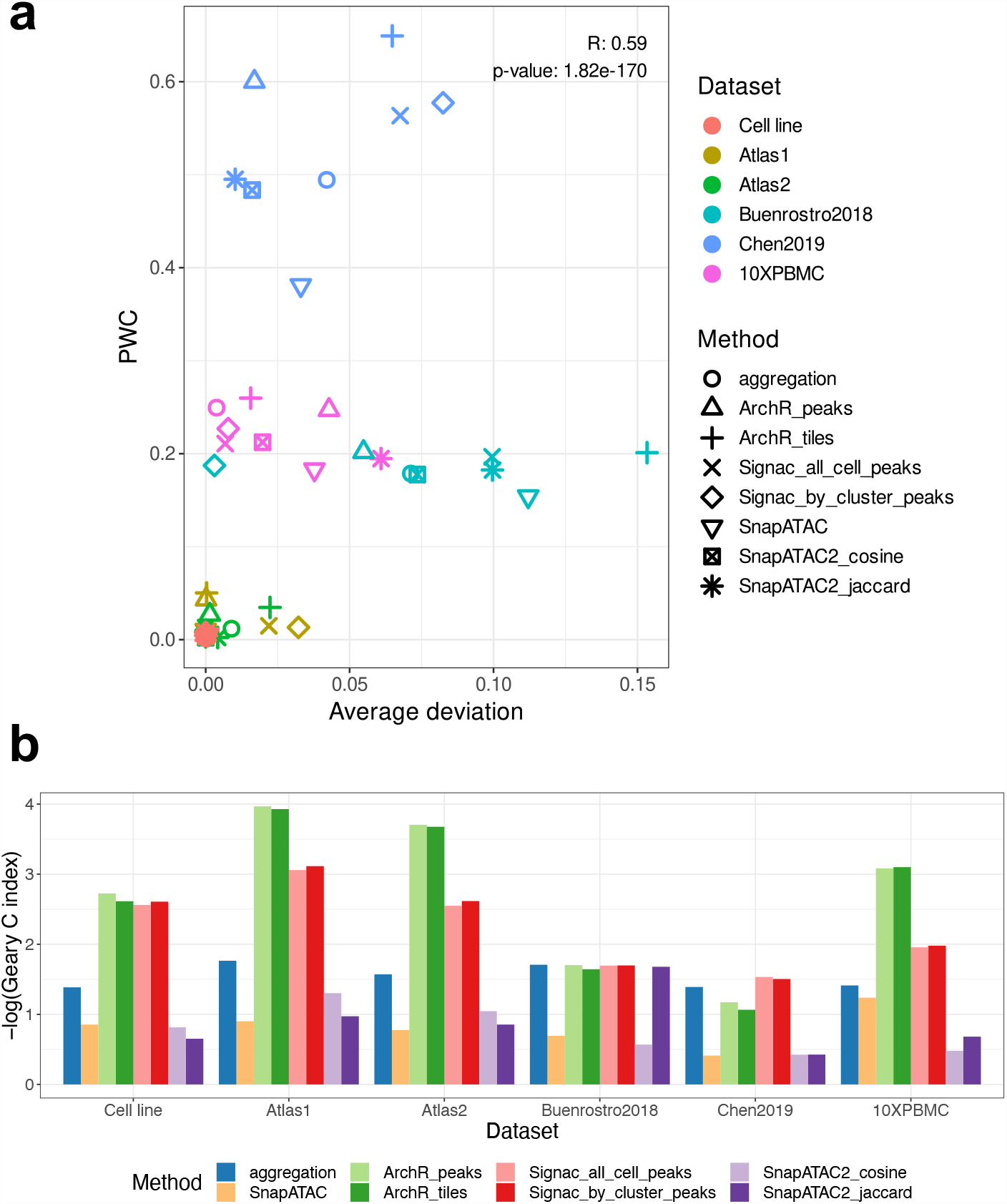
**a** Related to Figure S9, S10. x-axis is the deviation of ARI from 1, averaged across resolutions and random seeds. y-axis is the averaged PWC score of each dataset and method. **b** Spatial autocorrelation of library sizes measured by Geary’s C index.

**Figure S12:**
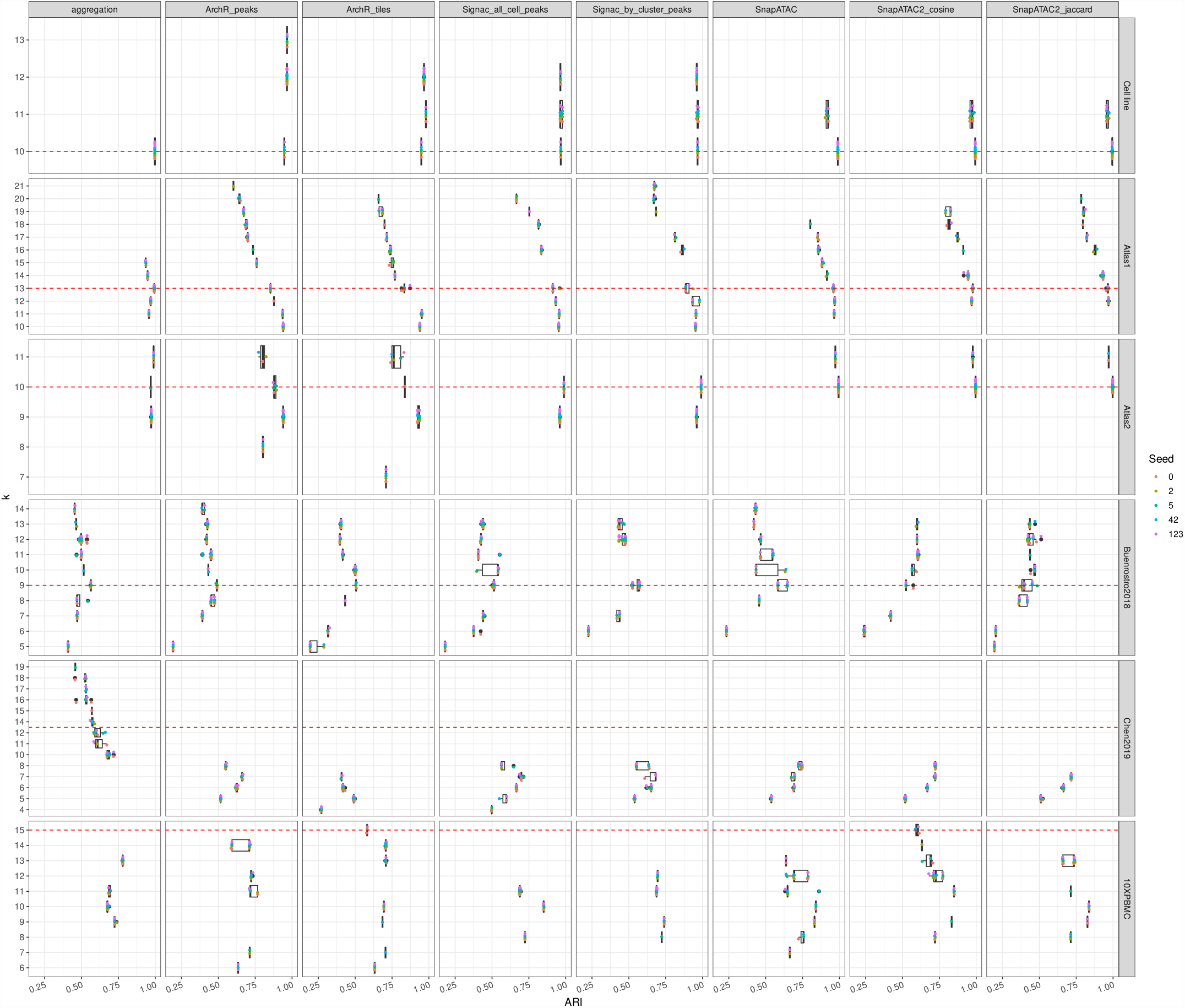
Boxplots of ARI between predicted clusterings and the true cell types, across datasets, methods, and the predicted number of clusters. The red horizontal lines are the ground truth cluster numbers.

**Figure S13:**
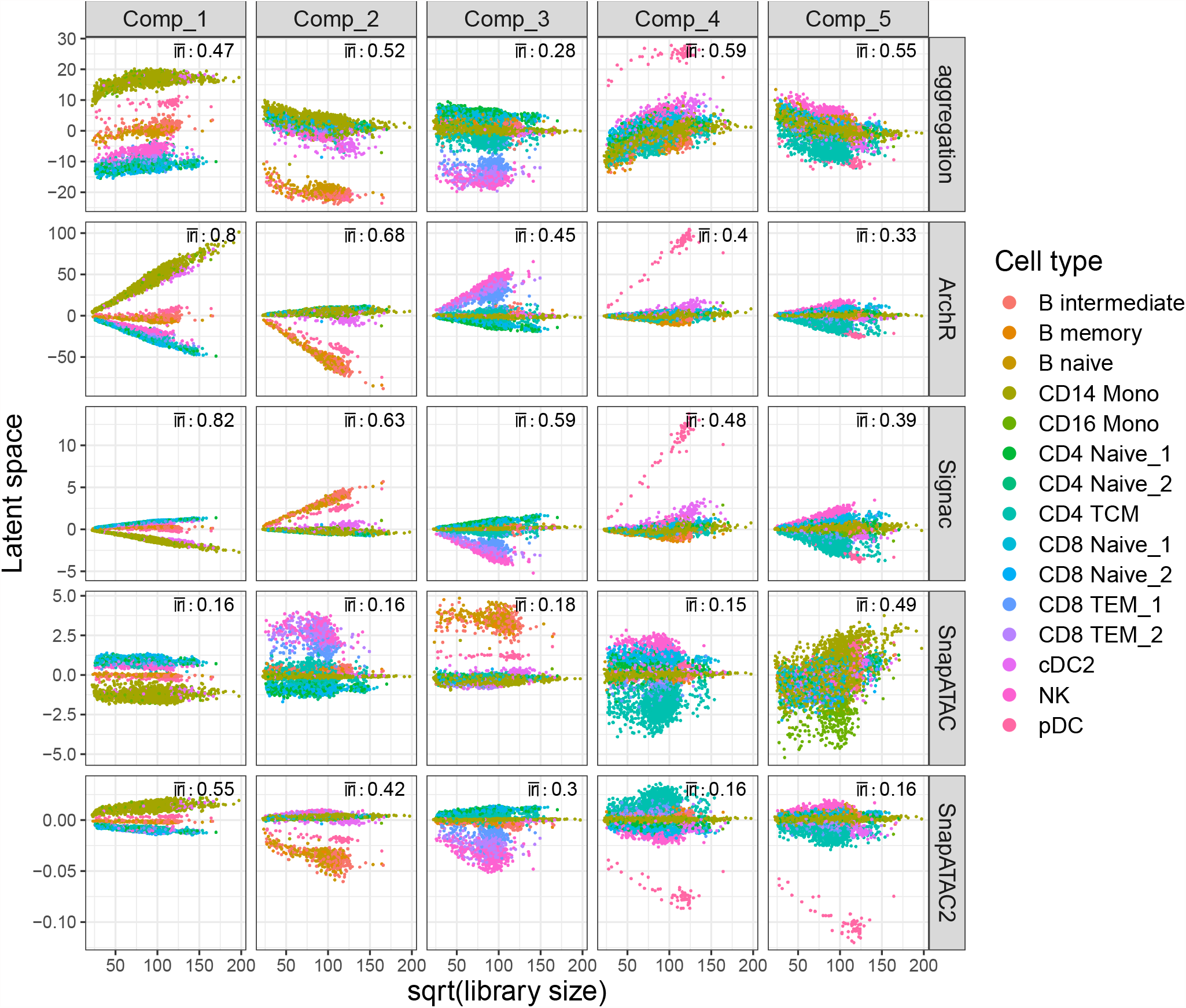
Scatter plots of the latent component value against the square root of fragment counts for dataset 10XPBMC. Colors are cell types, and the absolute Pearson’s correlation coefficient between x- and y-axis are calculated and averaged across cell types for each latent components.

**Figure S14:**
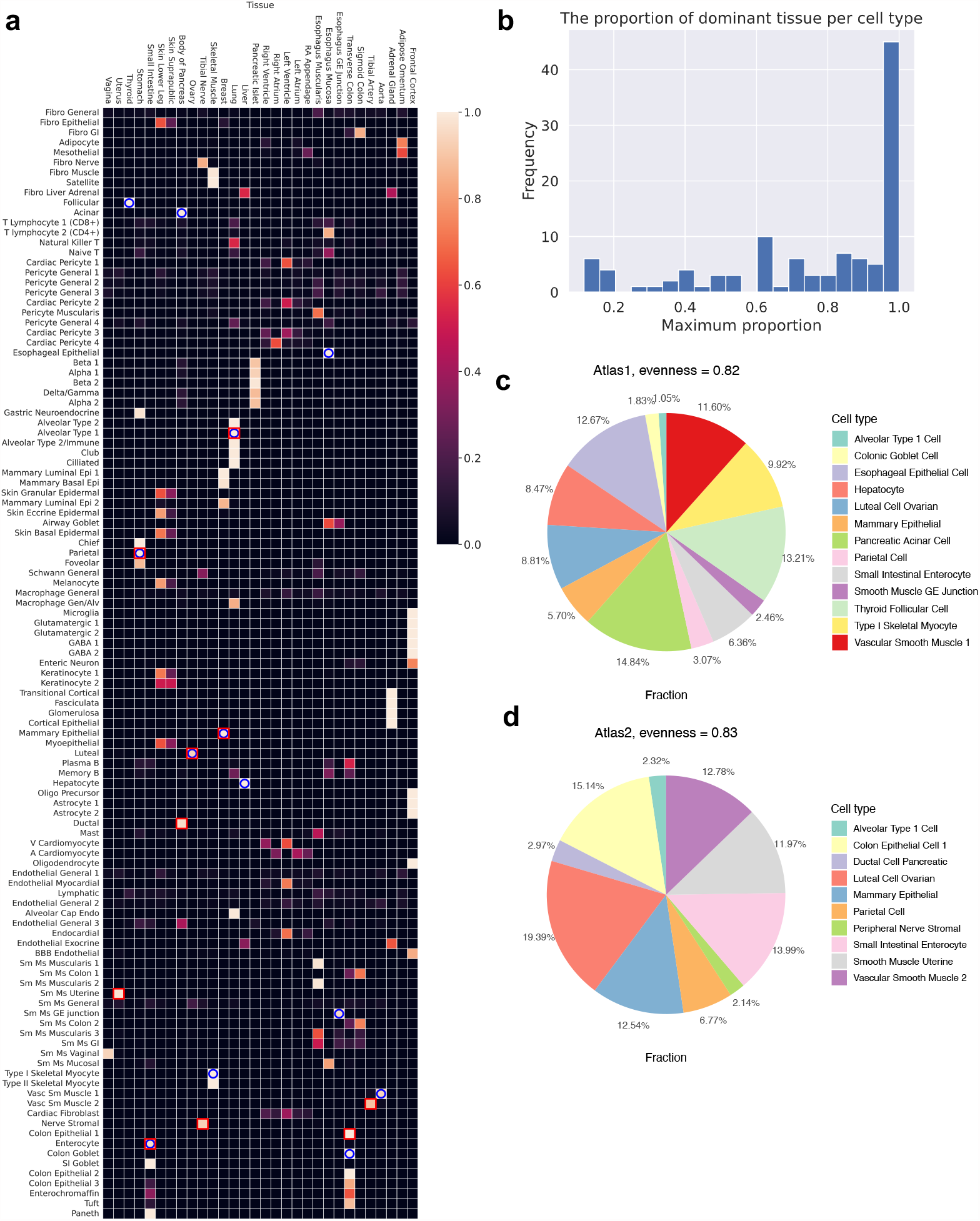
**a** Tissues and cell types in the human adult scATAC-seq atlas, with colors corresponding to column-wise proportion. Blue circles highlight the selected cell classes in Atlas1, and red rectangles highlight cell types in Atlas2. **b** Histogram of the proportion of dominant tissue for each cell type. For most cell types, the most dominant tissue contributes more than 85% cells. **c,d** Cell types and the proportion of our datasets Atlas1, Atlas2. Cell type evenness is calculated.

**Figure S15:**
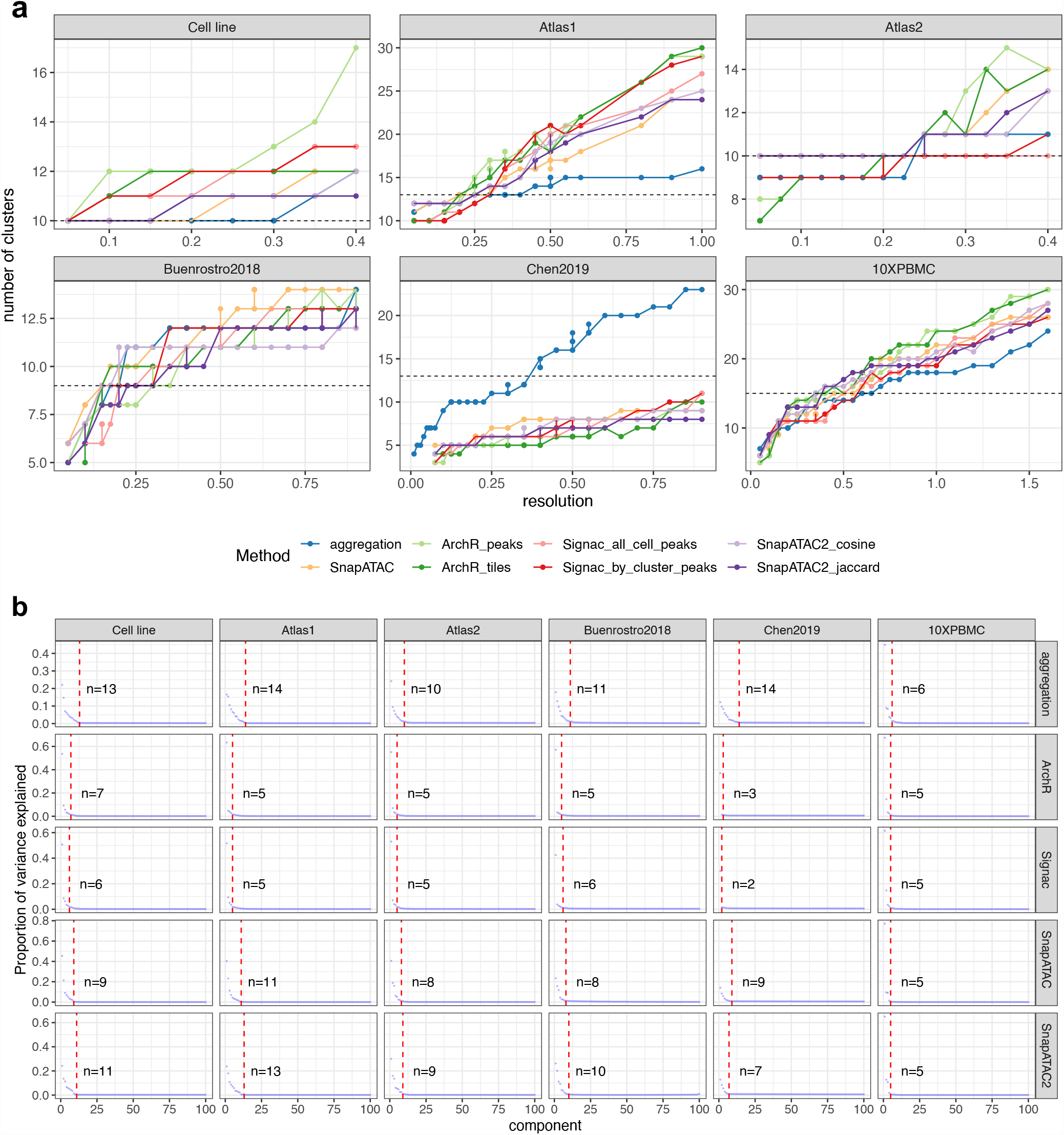
**a** The range of resolutions searched for each dataset, and how the number of clusters change as resolution changes. **b** Elbow plots for each method and datasets.

